# Ecological constraints to mirror life

**DOI:** 10.64898/2026.05.07.723461

**Authors:** Jordi Pla-Mauri, Victor Maull, Andrea Tabi, Yelyzaveta Shpilkina, Victor de Lorenzo, Ricard Solé

## Abstract

Our biosphere exhibits remarkable diversity yet is constrained by universal organizational principles, including molecular homochirality. Advances in synthetic biology have raised the possibility of engineering alternative life forms based on mirror-image biomolecules, prompting both technological interest and biosecurity concerns. While current discussions of mirror life largely emphasize molecular feasibility and cellular function, its potential establishment in natural environments remains poorly understood. Here, we develop a theoretical framework to assess the invasion potential of mirror organisms within existing ecosystems. Using population-level models that incorporate resource competition, metabolic constraints, and ecological network interactions, we show that mirror life might face severe limitations arising from both nutrient incompatibility and competitive exclusion by established biota. In particular, the reliance on rare or achiral substrates and the asymmetry of interactions with natural organisms constrain growth and persistence across a broad range of ecological conditions. These results highlight the importance of ecological constraints in evaluating the risks and feasibility of synthetic life.

## INTRODUCTION

The biosphere is a complex adaptive system, comprising an immense diversity of life forms that occupy habitats spanning a vast range of environmental conditions, from stable and nutrient-rich habitats to extreme environments. Despite this remarkable diversity, life on Earth exhibits a number of striking regularities that suggest the presence of universal constraints shaping biological organization across scales [1, 2]. Among these constraints is molecular chirality. Naturally occurring biological polymers are composed almost exclusively of one enantiomeric form of their constituent monomers, a property known as homochirality. Its origins have been extensively studied in the context of the origin of life, where it has been interpreted as the result of nonequilibrium symmetry-breaking processes in prebiotic chemical systems [3–6]. Once established, homochirality became a defining feature of biological chemistry, deeply embedded in the structure and function of living systems [7].

Can alternative life forms, based on different molecular principles, be created in the lab? In recent decades, synthetic biology has advanced rapidly toward the engineering of living systems, including the modification and expansion of the genetic code itself. Experimental efforts have demonstrated the possibility of incorporating non-canonical amino acids, redesigning codon assignments, and engineering orthogonal ribosomes capable of decoding alternative genetic instructions [8, 9]. These developments promise major advances in biotechnology and biomedicine, including novel therapeutics, programmable cells, and synthetic metabolic pathways [10, 11]. Its importance goes beyond circuit and cellular engineering, and it includes crops [12] and natural ecosystems [13–16].

However, the engineering of life raises significant biosecurity and ecological concerns, particularly regarding the release of engineered organisms into open environments. Such concerns are longstanding. During the early development of recombinant DNA technologies, debates over potential risks led to precautionary guidelines and temporary moratoria [17, 18]. More recently, attention has focused on “mirror life”: hypothetical organisms built from biomolecules of opposite chirality to those of extant life [19, 20]. Although current science remains far from creating such organisms—given the immense conceptual and technical challenges, and the absence of synthetic systems approaching even minimal cellular complexity [19, 21, 22]—mirror molecular biology has already revealed diverse potential applications. These include therapeutics, drug delivery systems, biosensors, plastic-degrading enzymes, and insights into biomolecular structure, homochirality, and the origin of life [21–27]. Many of these advantages stem from the high biostability of mirror compounds and their reduced immune recognition [22, 23]. These same properties, however, also raise safety concerns reminiscent of “gray goo” scenarios [28]. Because mirror organisms would likely resist natural pathogens and evade existing ecological controls, their accidental release could pose novel biosecurity risks—even with containment strategies such as synthetic auxotrophy [22]. In particular, their interactions with natural life may be strongly asymmetric, while remaining largely undetected by established biological regulation. Is that the case?

Most current discussions of mirror life risks are grounded in molecular and cellular considerations, emphasizing biochemical compatibility, pathogenicity, and metabolic autonomy [19, 20, 22]. However, a rigorous and constructive debate regarding biosecurity cannot proceed solely on these reductionist premises. Whether a novel life form could successfully invade and persist within the extant biosphere ultimately depends on ecological and evolutionary dynamics operating at population and ecosystem levels. Any assessment that neglects this ecological scale and the stringent constraints imposed by established environmental networks remains fundamentally incomplete and potentially misleading. Notably, any mirror organism released into the natural environment would face a severe nutrient limitation because most natural carbon sources would be unusable due to the opposite chirality. Their metabolism would be restricted to rare or specialized substrates, unless engineered to rely on achiral feedstocks [22]. Though, some argue that since many bacteria can grow in growth media without chiral nutrients, mirror organisms would be able to do the same, and that the amount of required achiral compounds would be enough to sustain mirror life [19].

Any alternative biological machinery would need to compete with highly optimized organisms embedded in dense, coevolved ecological networks. In this context, it should be noted that, despite longstanding speculation about the possibility of a “shadow biosphere”, i.e., a hypothetical coexisting life form with distinct molecular foundations, no empirical evidence for such alternative life has been found [29–31]. The absence of parallel molecular ecologies suggests that strong ecological and evolutionary constraints may prevent the long-term co-existence of fundamentally different biochemical systems. This empirical “silence” underscores that meaningful discourse on synthetic life must be anchored in an understanding of these environmental barriers, rather than hypothetical molecular capabilities alone.

In this paper, we develop a theoretical framework to evaluate the invasion potential of mirror life within existing ecosystems. Using different population-level models, we show that the structure and dynamics of the current biosphere can place stringent restrictions on their ability to establish and propagate. These limitations arise from the nonlinear dynamics of ecological communities, where resource competition, trophic interactions, and network-level organization determine the conditions for persistence. It is important to note that all models and scenarios described below treat mirror organisms as highly competitive invaders with unlimited mirror resource availability and the same biological efficiency and competitiveness as the extant organisms, despite real-world conditions that favor the established life forms (Table I). Our results indicate that, beyond engineering feasibility, these systemic ecological factors might strongly limit the viability of mirror organisms.

**Table 1.** Summary of the ecological models used to assess the potential spread of mirror life. Despite spanning closed, open, multispecies, and host-associated systems, all models consistently indicate that mirror life faces strong ecological constraints—primarily due to resource competition, metabolic asymmetries, and environmental filtering—making its successful invasion and persistence highly unlikely under realistic conditions.

| Model | Ecological Setting | Model Assumptions | Real-world conditions |
| --- | --- | --- | --- |
| Closed ecosystem model | Closed mass-conserving ecosystem | No external fluxes; strict mass conservation; organisms require fixed stoichiometric combination of achiral and chiral resources; nutrients are fully recycled; cross-feeding inefficient | Assumes unlimited access to mirror-compatible substrates, identical metabolic efficiencies between mirror and natural organisms, and neglects energetic costs of mirror biosynthesis. |
| Chemostat model | Open continuous-flow system | Constant inflow and dilution; achiral and chiral substrates externally supplied; dilution acts as mortality | Assumes persistent external supply of mirror-compatible resources and homogeneous mixing, while neglecting realistic scarcity of abiotic feedstocks, biofilm formation, and competitive specialization of extant microbes. |
| Predator-mediated closed model | Closed ecosystem with top-down regulation | Natural organisms controlled by predator/phage population; mass conservation and full recycling maintained | Assumes predators exclusively target natural organisms and ignores the possibility that mirror organisms would also suffer mortality from generalized grazing, abiotic stress, toxins, or partial immune recognition. |
| Multispecies ecosystem model | Diverse microbial community | Many resident species compete for substitutable resources; mirror variants have reduced access to chiral substrates | Assumes mirror variants retain nearly all ecological functionality of their natural counterparts despite lacking access to many chiral metabolic pathways; ignores coevolved mutualisms and microbiome-level exclusion effects. |
| Immune-response model | Within-host infection dynamics | Innate immune system partially recognizes mirror bacteria; macrophage-like predator-prey dynamics | Assumes only partial immune recognition and neglects complementary defenses such as adaptive immunity, complement cascades, mucosal barriers, and ecological competition with resident microbiota. |

## METHODS

Understanding whether mirror life could successfully spread within existing ecosystems requires moving beyond molecular and cellular considerations toward an explicit treatment of population dynamics. In particular, the establishment of a novel biological entity depends on its ability to grow when rare, compete for limiting resources, and persist within complex ecological networks. These questions are naturally framed within the mathematical theory of biological invasions, which provides a general framework to analyze the conditions under which a new species can invade and spread in a resident community [32–35]. Invasion dynamics emphasize key processes such as resource competition, demographic growth, spatial spread, and interaction structure, all of which determine whether an initially rare population can overcome extinction and establish itself. Within this context, mirror life can be treated as an invading lineage whose success depends not only on its intrinsic growth capabilities but also on its ecological compatibility with the extant biosphere. This perspective motivates the use of mathematical models that explicitly describe the coupled dynamics of populations and resources, allowing us to assess the conditions under which mirror organisms may fail or succeed in spreading.

We consider the dynamics of a resident ecological community interacting with a potential mirror-life lineage through a shared pool of molecular resources. The resident biosphere is represented by *K* populations

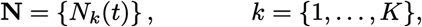

and the mirror lineage by a population *M* (*t*). Resources are partitioned into three classes: natural chiral compounds *C*_*n*_, mirror-chiral compounds *C*_*m*_, and achiral compounds *A*. These classes differ in their biochemical accessibility and mediate indirect interactions between populations. A general model reads:

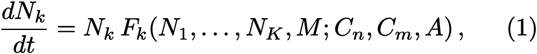

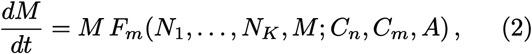

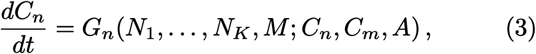

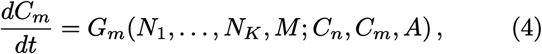

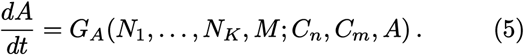

with *k* = 1, …, *K*. Here, *F*_*k*_ and *F*_*m*_ represent effective per-capita growth rates, incorporating resource uptake, metabolic constraints, and ecological interactions (e.g., competition, facilitation, or trophic effects). The functions *G*_*i*_ describe the dynamics of resource pools, including supply, depletion, transformation, and recycling. A key feature of the model is the asymmetric accessibility of chiral resources. Natural populations primarily exploit *C*_*n*_, while mirror populations primarily exploit *C*_*m*_, with both potentially accessing the achiral pool *A*. This asymmetry is encoded implicitly in the functional dependence of *F*_*k*_ and *F*_*m*_ on resource variables, without specifying particular uptake forms. As a consequence, interactions between natural and mirror populations are mediated indirectly through shared resources (notably *A*) and through modifications of the resource landscape. The framework accommodates both closed and open ecological settings. Open systems (e.g., chemostats or environmental fluxes) are described by allowing *G*_*i*_ to include external input and loss terms, so that no global conservation law applies.

To assess the potential for mirror life to establish, we consider the dynamics of *M* when rare. Linearizing around a resident equilibrium 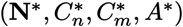 with *M* = 0, the invasion growth rate is given by

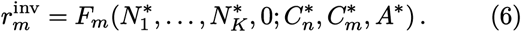

Mirror life can invade only if 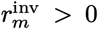. Otherwise, it declines when rare and cannot establish. This quantity provides a unifying criterion that links molecular constraints (resource accessibility) with ecological interactions (community structure and competition). The specific models analyzed in the Results section correspond to particular choices of the functions *F*_*k*_, *F*_*m*_, and *G*_*i*_, representing different ecological scenarios (e.g., closed systems, chemostats, and network-mediated interactions). This general formulation provides a common mathematical framework for comparing these cases.

## RESULTS

To assess the ecological feasibility of mirror life, we develop minimal population-dynamical models of native and mirror replicators competing for shared chiral and achiral resources. Across scenarios—from closed systems to multispecies communities—we isolate the key drivers of invasion and persistence. In all cases, resource competition and the presence of established communities strongly limit the spread of mirror life.

### Mirror life in a closed ecosystem

Our first case study involves a closed ecosystem. This class of model ecosystems, which includes closed eco-spheres and life-support systems [36], is well known in population biology [37, 38] and has received renewed attention in recent years within the context of synthetic biology [15, 39–41]. We consider a closed ecosystem comprising natural (*N*) and mirrored (*M*) chiral organisms that compete for achiral (*A*) and homochiral resources (*C*_*n*_, *C*_*m*_). The system is governed by strict mass conservation, characterized by a specific metabolic constraint: while organisms may assimilate achiral or chiral resources independently from the environment, internal biomass synthesis requires these components to be combined in a fixed stoichiometric ratio of *λ* : (1*− λ*). Upon mortality at a rate *δ*, organisms fully recycle their mass back into the resource pool according to this fixed ratio. A diagram of the interactions is provided in Fig. 1b.

**FIG. 1.**
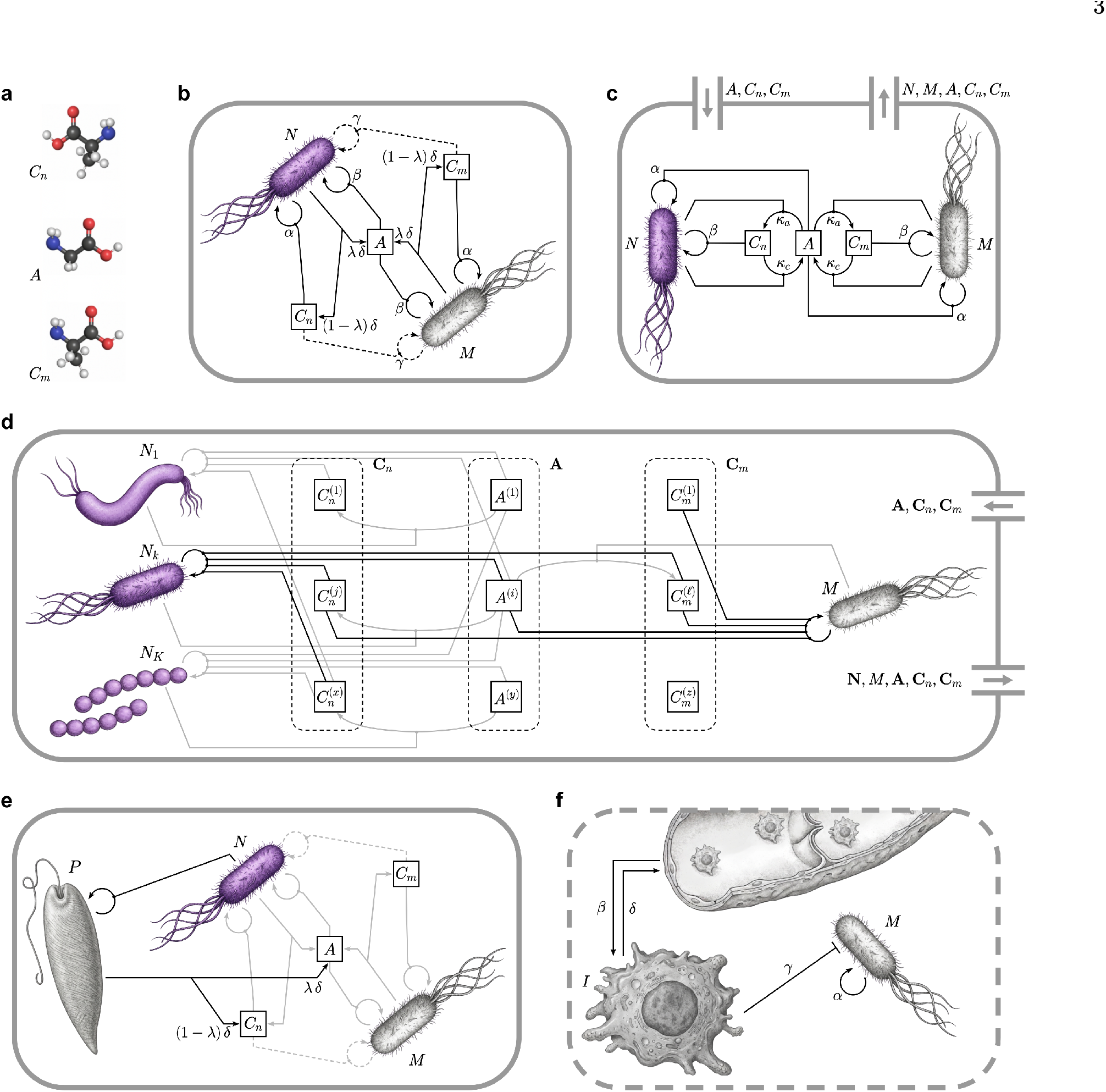
Ecological interactions between natural and mirror life. (a) Three classes of molecular resources are considered: standard chiral metabolites compatible with extant biochemistry (*C*_*n*_), their mirror-chiral counterparts (*C*_*m*_), and achiral compounds (*A*), which can in principle be accessed by both types of organisms. (b) Closed ecosystem. Two microbial populations are present: a resident community of natural organisms (*N*) and a mirror-life population (*M*). Both populations transform and recycle resources through metabolic processes and contribute to the production of shared and chiral compounds, with a fraction of flux channeled into achiral resources and the remainder into chiral pools. Interactions between *N* and *M* are indirect and mediated by resource availability, with no external inputs or losses. (c) Controlled open system (chemostat-like dynamics). Resources (*A, C*_*n*_, *C*_*m*_) are supplied through external fluxes, and both resources and populations are subject to dilution. Resource uptake and conversion are characterized by effective metabolic rates, while growth depends on resource-dependent feedbacks. In this regime, competition between natural and mirror populations is mediated by shared access to achiral compounds and by the transformation of resource pools under continuous flow conditions. (d) Generalized multispecies open system. Multiple resident populations (**N**) and a mirror variant of a resident (*M*) compete for externally supplied resources (**A, C**_*n*_, **C**_*m*_) under dilution. Primary interactions with resources are shown in black; all other interactions, including resource interconversions, are shown in grey. (e) Closed ecosystem with a natural predator. The system from (b) is extended by adding a predator population that targets the natural organisms. Interactions already present in the closed ecosystem are dimmed in grey. (f) Open system with an immune response. A mirror bacterial population grows in an open tissue environment and is subject to predation by macrophages. A vessel on top represents source for the arrival and removal of macrophages at defined rates.

**FIG. 2.**
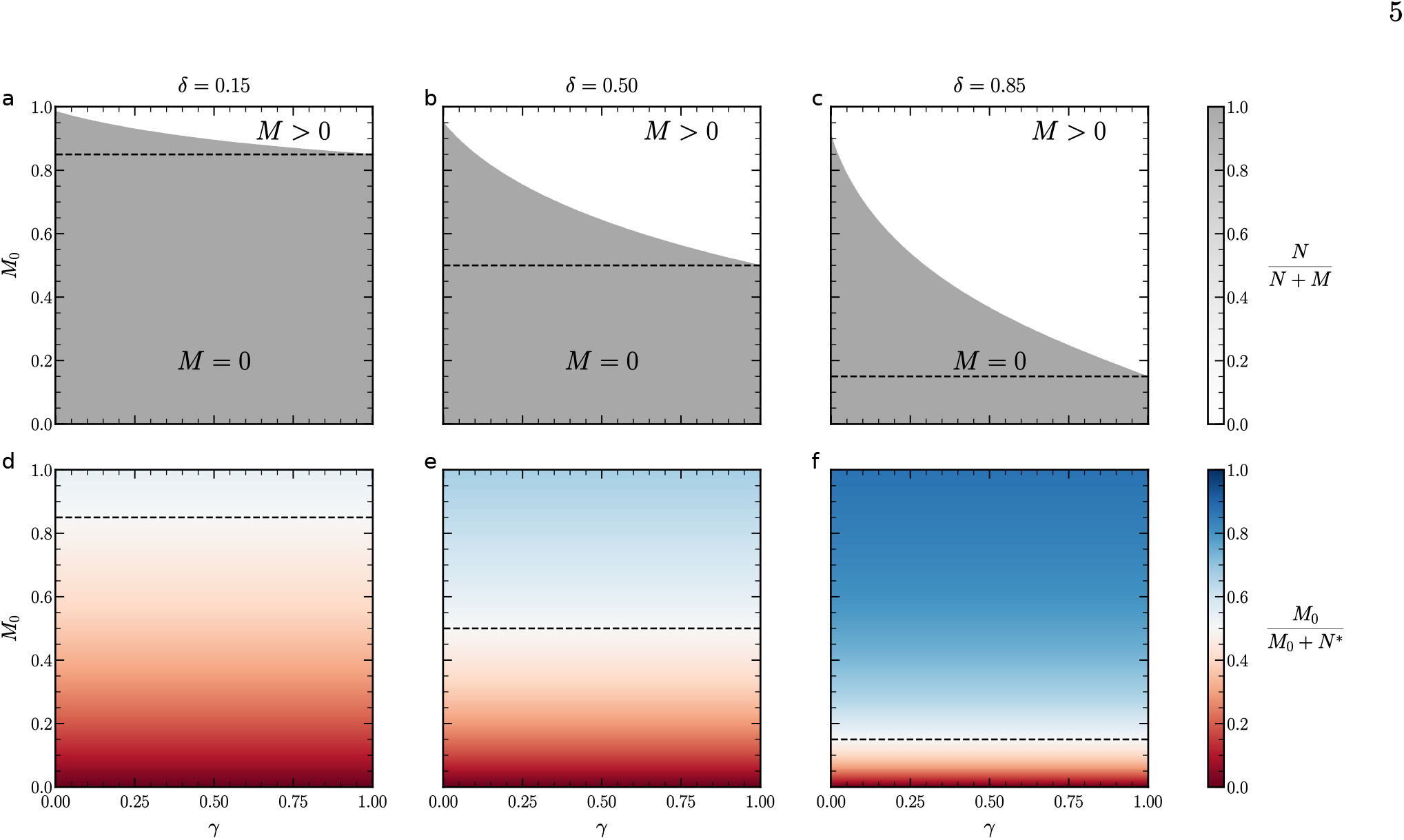
Steady state after invasion by mirror inoculum in a closed ecosystem. Panels (a), (b), and (c) show the biomass fraction *N/*(*N* + *M*) for *δ* = 0.15, *δ* = 0.5, and *δ* = 0.85, respectively. Panels (d), (e), and (f) show the corresponding resource fraction *C*_*n*_*/*(*C*_*n*_ + *C*_*m*_). Dashed lines mark the manifold where the initial mirror inoculum *M*_0_ equals *N*\*, the steady-state biomass of the natural population in the absence of the mirror organism. Mirror invasion is only possible when *M*_0_ *> N*\*—effectively requiring an initial mirror inoculum that supplies *at least as much biomass as the established living natural population*—but this condition is necessary, not sufficient, for mirror survival and displacement of *N*, and it approaches equality only in cases where cross-feeding efficiency is high (*γ ≈ β*). As the death rate δ increases, more resources become available in the closed system, progressively expanding the conditions under which the mirror population can successfully invade. Parameters: *α* = 1, *β* = 1, *λ* = 0.1.

**FIG. 3.**
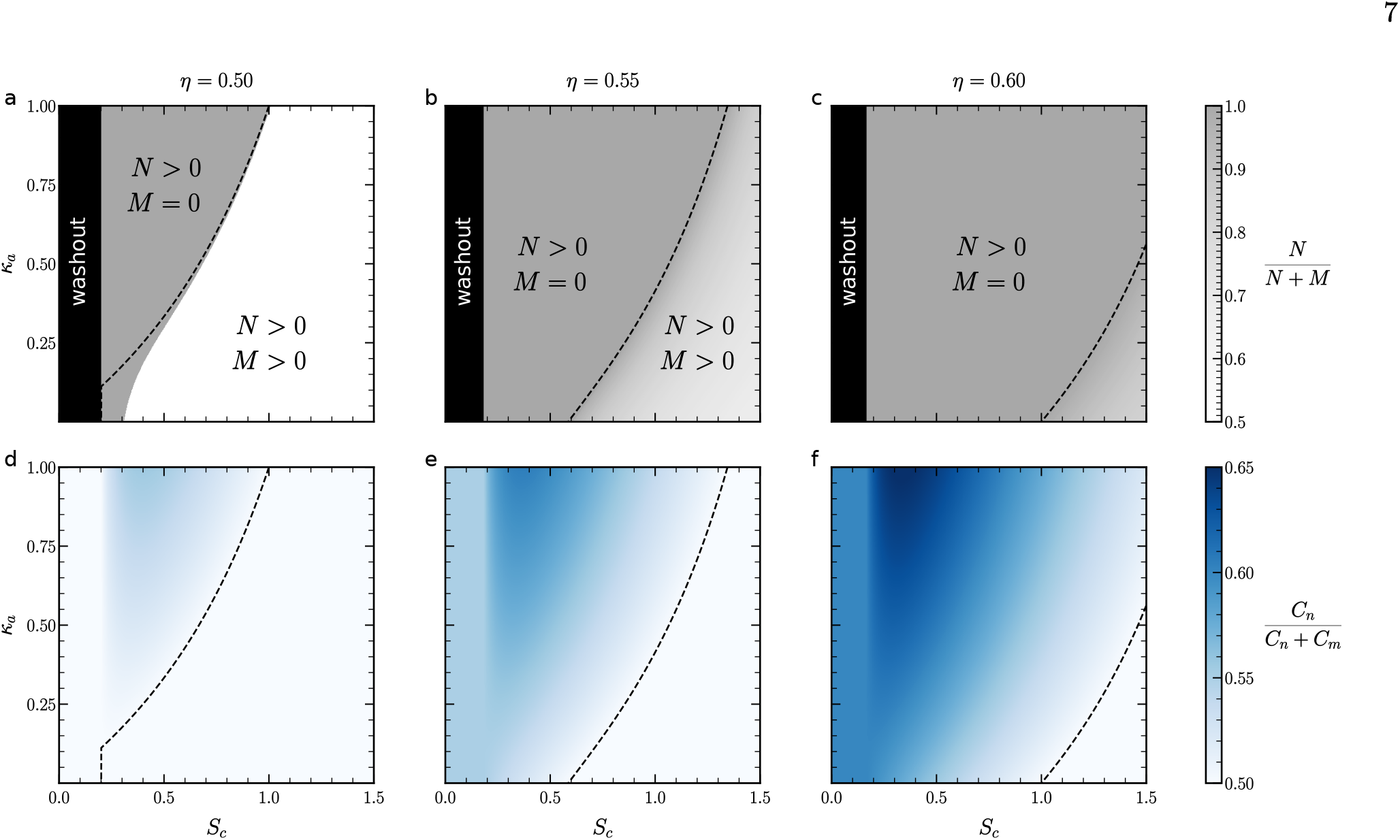
Steady state after invasion by mirror inoculum in a bioreactor. Panels (a), (b), and (c) show the biomass fraction *N/*(*N* + *M*) for *η* = 0.5, *η* = 0.55, and *η* = 0.6, respectively. Panels (d), (e), and (f) display the corresponding resource fraction *C*_*n*_*/*(*C*_*n*_ + *C*_*m*_). At low supplied nutrient concentrations (*S*_*c*_), growth cannot outpace dilution, leading to complete washout of all organisms. As the nutrient supply increases, the natural population establishes a steady state. If it can construct a niche for itself (*κ*_*a*_ *>* 0), it becomes more resilient to mirror invasions, and progressively larger values of *κ*_*a*_ broaden the region of parameter space where the mirror population is completely excluded. At very high resource supply rates, competition weakens as resources cease to be limiting, allowing both populations to coexist. Parameters: *D* = 1, *µ* = 1, *γ* = 1, *κ*_*c*_ = 0.

To sustain a stable biotic steady state, we assume full resource recycling, which yields the following dynamical system

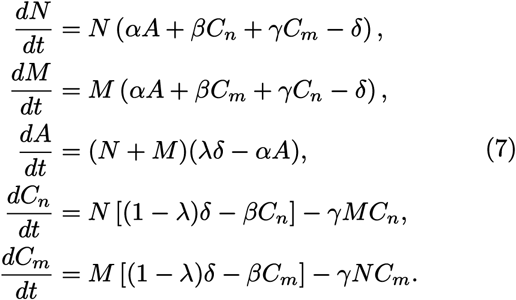

where *α*, quantifies the utilization efficiency of the common achiral resource, *β* quantifies the assimilation efficiency of the cognate chiral resource, and *γ* denotes the assimilation efficiency of the non-cognate (cross-fed) chiral resource. We assume that *γ < β* reflecting biological homochirality: utilizing non-cognate resources requires energetically costly racemization, rendering cross-feeding inherently less efficient than direct assimilation.

To ensure full resource recycling the consumption coefficients are set equal to the growth efficiency parameters. Let the total mass of the system be defined as Ω = *N* + *M* + *A* + *C*_*n*_ + *C*_*m*_. It can be checked that the time derivative of the total mass vanishes

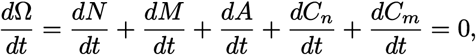

and consequently, total mass Ω is a preserved quantity.

Let us analyse now the steady state corresponding to the extinction of the mirrored population (*M*\* = 0) and the persistence of the natural population (*N*\* *>* 0), which should be a proxy of the current state in the biosphere. Given the assumption of *N*\* *>* 0 and *M*\* = 0,

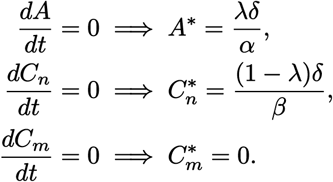

These values represent the critical resource concentrations required to balance growth against mortality. Since no mechanism exists to generate *C*_*m*_ in the absence of population *M*, the concentration 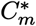 vanishes at equilib-rium. The sole exception occurs if cross-feeding is absent (*γ* = 0); in this limiting case, 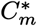 remains constant at its initial value.

The population equilibrium can then be derived by uti-lizing the mass conservation,

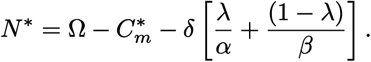

Therefore, a viable biotic steady state exists if and only if the active mass exceeds the minimum maintenance threshold defined by

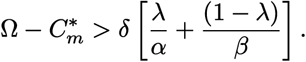

Should this condition fail to hold, the system collapses to a state of total extinction where *N*\* = *M*\* = 0.

The invasion fitness 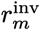, defined as the per capita growth rate of a rare invader (*M →*0), is obtained by evaluating the growth equation for *M* at the resident steady state:

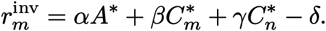

Substituting the steady state values and rearranging terms yields the general condition for invasion:

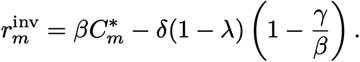

Thus, the mirrored population can successfully invade if and only if the residual supply of the mirrored resource exceeds the effective metabolic threshold:

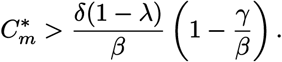

When cross-feeding is present (*γ >* 0), the resident population depletes mirror resources (*C*_*m*_), making invasion effectively impossible (given *γ < β*). Only in the limiting case *γ* = 0 can mirror resources persist, and even then invasion requires a sufficiently large initial supply. More generally, cross-feeding imposes a strong asymmetry that drives negative invasion fitness for mirror life.

This scenario suggests that the intrinsic metabolic inefficiency of cross-feeding on non-cognate chiral resources would place mirror life at a competitive disadvantage against native life in closed ecosystems. The only cases where mirror organisms could potentially outperform its native counterparts is if a sufficiently large initial inoculum is introduced, effectively adding a stock of mirror resources to the ecosystem and temporarily circumventing the cross-feeding asymmetry.

Two other relevant case studies within the closed ecosystem scenario are explored in the Supplementary Material: autrotrophs (SM I) and bacteriostatic antibiotics (SM II).

### Mirror life in a chemostat

In contrast to the closed ecosystem, we now consider an open system: a chemostat where continuous inflow of fresh resources and outflow of reactor contents frees the system from the strict mass conservation that characterized the previous case (see Fig. 1c). Here, the resource dynamics are governed not by recycling from dead biomass, but by the interplay between external supply and possibly metabolic interactions. The achiral substrate is supplied at concentration *S*_*a*_, while the chiral substrate enters as enantiomeric pairs with total concentration *S*_*c*_, partitioned according to an enantiomeric ratio *η ∈* [0, 1] such that 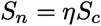 and *S*_*m*_ = (1 − *η*)*S*_*c*_, where *η* = 0.5 would represent an inorganic, racemic source. All resource types occupy the same metabolic niche and are the limiting factors for growth. The parameter *D* represents the dilution rate, which simultaneously regulates the influx of fresh resources and the washout of all reactor contents. Furthermore, the coefficients *κ*_*a*_ and *κ*_*c*_ quantify metabolic leakage rates, characterizing the conversion of achiral resources into chiral forms and vice versa relative to the uptake rates that directly drive population reproduction.

The resource dynamics are governed by the following system:

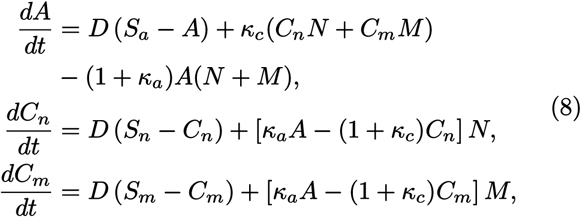

where the inflow concentrations of the normal and mirror chiral substrates, 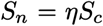 and *S*_*m*_ = (1 *−η*)*S*_*c*_ are defined to partition the total chiral supply *S*_*c*_ according to the enantiomeric ratio *η ∈* [0, 1]—a value of *η* = 0.5 would represent an inorganic source. The parameter *D* represents the dilution rate, which simultaneously regulates the influx of fresh resources and the washout of all reactor contents. Furthermore, the coefficients *κ*_*a*_ and *κ*_*c*_ quantify metabolic leakage rates, characterizing the conversion of achiral resources into chiral forms and vice versa relative to the uptake rates that directly drive population reproduction.

The biotic dynamics are governed by

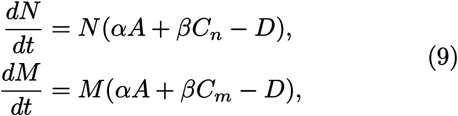

where *α* and *β* denote the conversion efficiencies of achiral (*A*) and cognate chiral (*C*_*n*_, *C*_*m*_) resources into biomass, respectively.

To determine the conditions for the existence of the normal-chiral population, we first analyze the system assuming the mirror-chiral population is absent (*M* = 0). Under this condition, the equation for the mirror-chiral substrate simplifies to

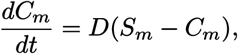

which at steady state yields the inflow concentration, 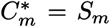. For the normal-chiral population to persist (*N*\* *>* 0), its net growth rate must be zero. This requires

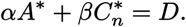

Similarly, the resource dynamics must be at equilibrium:

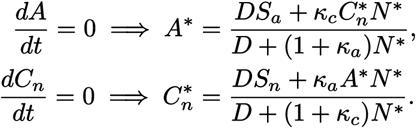

Substituting these algebraic solutions into the zero-net-growth condition for the population *N* yields a quadratic equation for steady-state biomass:

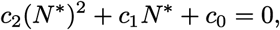

where the coefficients are determined by the system parameters:

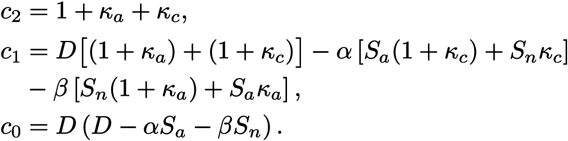

Given that all parameters that represent rates and concentrations must be positive, the quadratic coefficient *c*_2_ is strictly positive. The constant term *c*_0_ is negative if and only if the maximum potential growth rate exceeds the dilution rate, which constitutes the necessary condition for survival:

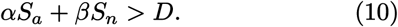

This guaranties exactly one positive real root and one negative real root.^1^ Since biomass cannot be negative, there exists a unique positive steady state *N*\* if and only if equation (10) is satisfied. The value of this steady state is given by

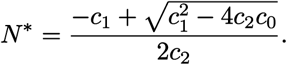

We now evaluate the stability of this mono-chiral steady state against the introduction of the mirror-chiral population. Introducing a small perturbation *M* → 0^+^, the invader will grow if its initial per capita growth rate is positive:

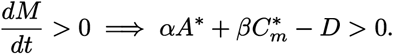

Given the previously derived steady state, this condition becomes

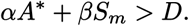

Substituting *D* in the invasion inequality with the zeronet-growth condition of the resident population *N*\* and simplifying yields

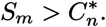

In the regime where consumption dominates leakage 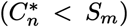, the specific resource niche of the invader is richer than that of the resident, allowing mutual invasibility and leading to stable coexistence. Conversely, if metabolic leakage is sufficiently high such that organisms generate their specific resource faster than it is depleted, or if the input ratio heavily favors the resident 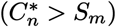, the resident maintains a resource advantage. Under these conditions, the mono-chiral state is stable against invasion, resulting in competitive exclusion.

### Mirror life with predators

To account for top-down regulatory mechanisms inherent in natural ecosystems, we extend the model to include a specialized predator (or phage) population, denoted by *P*, which targets the natural chiral organism *N*. We assume strict mass conservation wherein the predator converts consumed biomass into its own with unit efficiency. Upon mortality, the predator’s biomass is fully recycled into the achiral resource *A* and the cognate chiral resource *C*_*n*_ according to the same stoichiometric ratio as the other organisms.

The resulting dynamical system is governed by

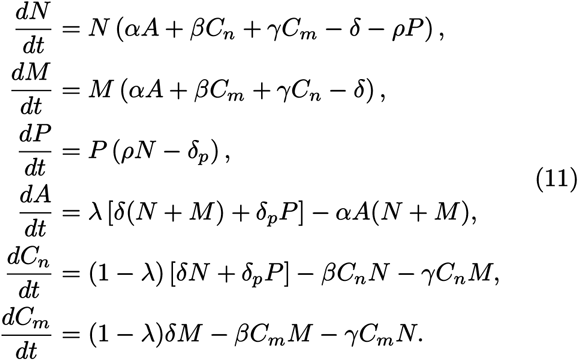

where *ρ* indicates predation rate. The total mass Ω = *N* + *M* + *P* + *A* + *C*_*n*_ + *C*_*m*_ remains a conserved quantity. We can analyze the steady state characterized by the extinction of the mirrored population (*M*\* = 0) and the coexistence of the natural population and its predator (*N*\* *>* 0, *P*\* *>* 0). For this predator-controlled system, equilibrium for the prey density is given by

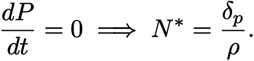

Substituting this into the resource balance equations, the equilibrium concentrations are found to be

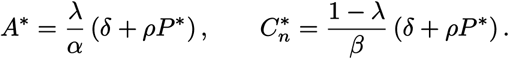

The steady state value of *P*\* is then obtained from mass conservation, explicitly accounting for this residual pool

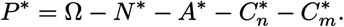

In the absence of the mirrored population (*M*\* = 0), the fate of 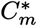 depends on the cross-feeding: if *γ* = 0, 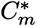 remains fixed at its initial concentration; conversely, if *γ >* 0, the resource will be fully depleted 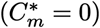.

The invasion fitness 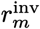 of a rare mirrored population is given by

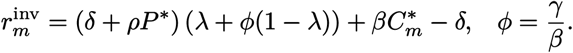

Successful invasion requires 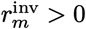, which results in the condition

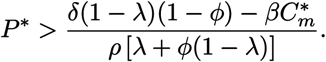

In this case, top-down control can facilitate invasion of the mirrored population by suppressing *N*\* and thus increasing residual resource concentrations 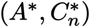. Additionally, any pre-existing reservoir of the specific chiral resource 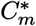 directly contributes to the invader’s growth potential, lowering the threshold of predator density required for successful establishment.

The addition of a predator that targets the natural chiral organism can facilitate invasion by a mirror organism by suppressing the natural population and freeing up exploitable resources. However, this effect is contextdependent: it applies primarily when the natural ecosystem is already degraded, or when the predator exerts unusually strong top-down control over the natural population despite the availability of resources that could otherwise support growth.

### Mirror life in a multispecies ecosystems

In the previous sections, we focused on simplified model ecosystems in which the natural community is represented by a single species. However, when assessing the potential impact of mirror life in real ecosystems, it is essential to consider diverse and interacting communities. We therefore consider a steady-state microbial ecosystem comprising *n* resident species competing for *m* substitutable resources. Unlike the simplest chemostat models, we allow each population *k* to experience a distinct loss rate *D*_*k*_. This loss rate may encapsulate dilution in a chemostat, density-independent mortality, predation, or viral lysis specific to that organism. At equilibrium, the resource vector **R**\* sustains the resident community such that, for each persisting species, its realized growth rate equals its own loss rate *D*_*k*_.

Select a resident organism *k* and construct a chiral mirror variant *k*′. This variant is metabolically identical to *k* regarding achiral resources but differs in its capacity to utilize chiral resources. While *k*′ consumes achiral resources identically to *k*, utilization of chiral resources necessitates specific racemases to convert environmental enantiomers into a usable form. If the variant lacks the requisite racemase for a specific chiral resource, or if the metabolic cost of expressing such enzymes is prohibitive, uptake is hindered. Consequently, for any resource vector, the growth rate of the mirror variant is bounded by that of the natural organism, with strict inequality arising whenever a chiral resource contributes to growth and remains inaccessible to the variant. The loss rate experienced by the mirror variant, 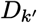, may differ from the resident’s *D*_*k*_ due to factors such as escape from specialized predators or phages that recognize chiral surface structures, or resistance to antibiotics targeting the natural enantiomer.

At the resident steady state **R**\*, the natural organism satisfies the equilibrium condition where its growth rate equals its own loss rate *D*_*k*_. The invasion fitness of the mirror variant is determined by its growth rate at **R**\* relative to its own loss rate 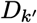. Unless the variant possesses the full complement of enzymes and racemases needed to metabolize every chiral resource without any associated fitness cost—an exceedingly unlikely scenario for complex resources—its metabolic capabilities form a strict subset of the resident’s, and its growth rate at equilibrium cannot surpass that of the resident. Invasion is thus possible only if the reduction in the mirror loss rate overcomes the metabolic deficit in chiral resource utilization.

This principle can be formalized using a simple chemostat-like system where the dynamics of resources **R** and species **X** are governed by

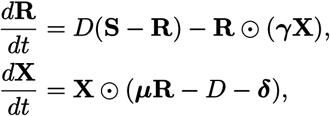

with **S** representing the inflowing resource concentrations, ***γ*** the yield coefficients for growth and resource interconversion, ***µ*** the matrix of per-capita growth rates per unit resource, *D* the chemostat dilution rate, and ***δ*** the population-specific losses due to microbial warfare, predation, and other ecological interactions.

To incorporate chiral constraints, the resource vector is divided into achiral (**A**), natural chiral (**C**_**n**_), and mirror chiral (**C**_**m**_) components. The growth rate of a resident species *k* is the sum of contributions from all three sets

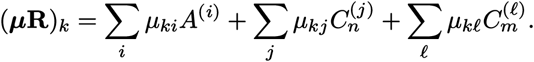

For its mirror variant *k*′, the uptake coefficients for achiral resources can be assumed to remain the same since achiral molecules interact identically with enzymes of either handedness 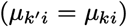. For chiral resources, however, the coefficients are bounded by the corresponding rates of the original strain: the mirror variant’s uptake of natural chiral resources is capped by the natural rates 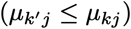, while its uptake of mirror resources cannot exceed the natural strain’s baseline ability to use them 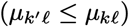. These constraints capture the possible lack of specific enzymes, racemases or the energetic cost of enantiomerization in both directions.

The invasion fitness 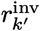 of the mirror variant is defined as its net growth rate at the resident steady state

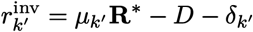

Although mirror resources appear in principle, they are typically rare in natural settings and can be neglected. Assuming 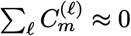, the resident satisfies

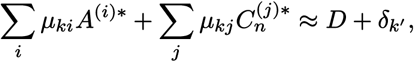

and since achiral contributions cancel out due to the previous assumption of equal coefficients, the invasion fitness simplifies to

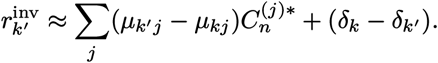

Because 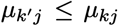 for all natural chiral resources, the first term is non-positive. Equality holds only in the idealized case where the variant matches the resident’s chiral uptake perfectly without cost. In practice, some chiral resource will either be inaccessible to the mirror variant or too costly to be efficiently exploited, so the term can be assumed strictly negative, establishing a chiral barrier to invasion. In the SM we analyze the invasion dynamics of ML invasion under open (SM III) and closed (SM IV) ecosystem frameworks.

Thus, in a resource-limited complex ecosystem where higher-order interactions are mediated through resources, the establishment of a mirror organism is highly constrained. Only under top-down control, where additional pressures, such as antibiotics or predation, play a significant role, can this chiral barrier be overcome (see also SM II). In such scenarios, the suppressed resident leaves exploitable resources in the environment, opening a niche that the mirror variant can occupy and potentially allowing the invader to succeed.

### Immune responses to mirror life

All previous models rely on an ecological framework and can be analyzed via invasion dynamics of the mirror strain. However, a distinct concern is that mirror life may evade immune recognition—e.g., macrophage detection—due to reversed molecular chirality, potentially enabling uncontrolled persistence. Although this is outside the ecological setting, it can be mathematically addressed considering the dynamics within the host [42–44]. Despite the complexity of immune responses [45], the core interaction can be captured by a minimal two-dimensional system between immune defense *I* (e.g., macrophages) and mirror bacteria *M* :

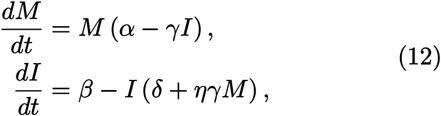

where *α* is the growth rate of the mirror cells, *γ* is the phagocytic clearance rate, *β* the recruitment rate (arrival rate) of new macrophages into the system, *η* the exhaustion index, representing the fractional loss of the macrophage population per successful predation event, and *δ* is the macrophage clearance rate.

There is a transcritical bifurcation at the parameter threshold *βγ* = *αδ* (Fig. 5a). In the cases where *βγ < αδ*, there exists a unique (unstable) fixed point with *M*\* = 0, *I*\* = *β/δ*. For any initial inoculum *M*_0_ *>* 0, the growth of mirror cells exceeds the maximum clearance capacity of macrophages. The system thus evolves towards unbounded growth of the mirror population (*M* → ∞). Instead, if *γ > γ*_*c*_ = *αδ/β*, the extinction point becomes locally asymptotically stable, and an additional fixed point enters the physical quadrant,

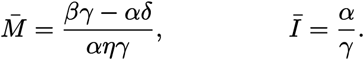

This saddle point acts as a separatrix between two divergent basins of attraction, creating a strong Allee effect (Fig. 5b). The small initial inocula 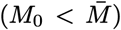 are effectively suppressed by the baseline macrophage population, leading to the stable (healthy) attractor (*M* →0). Large inoculum events 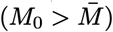 can still trigger a rapid exhaustion of the macrophage pool that outpaces the recruitment rate. In this case, if the density of transient macrophages falls below the critical threshold *I*\* = *α/γ*, the predation pressure becomes insufficient to counteract the growth of mirror cells, resulting in systemic invasion (*M*→ ∞).

**FIG. 4.**
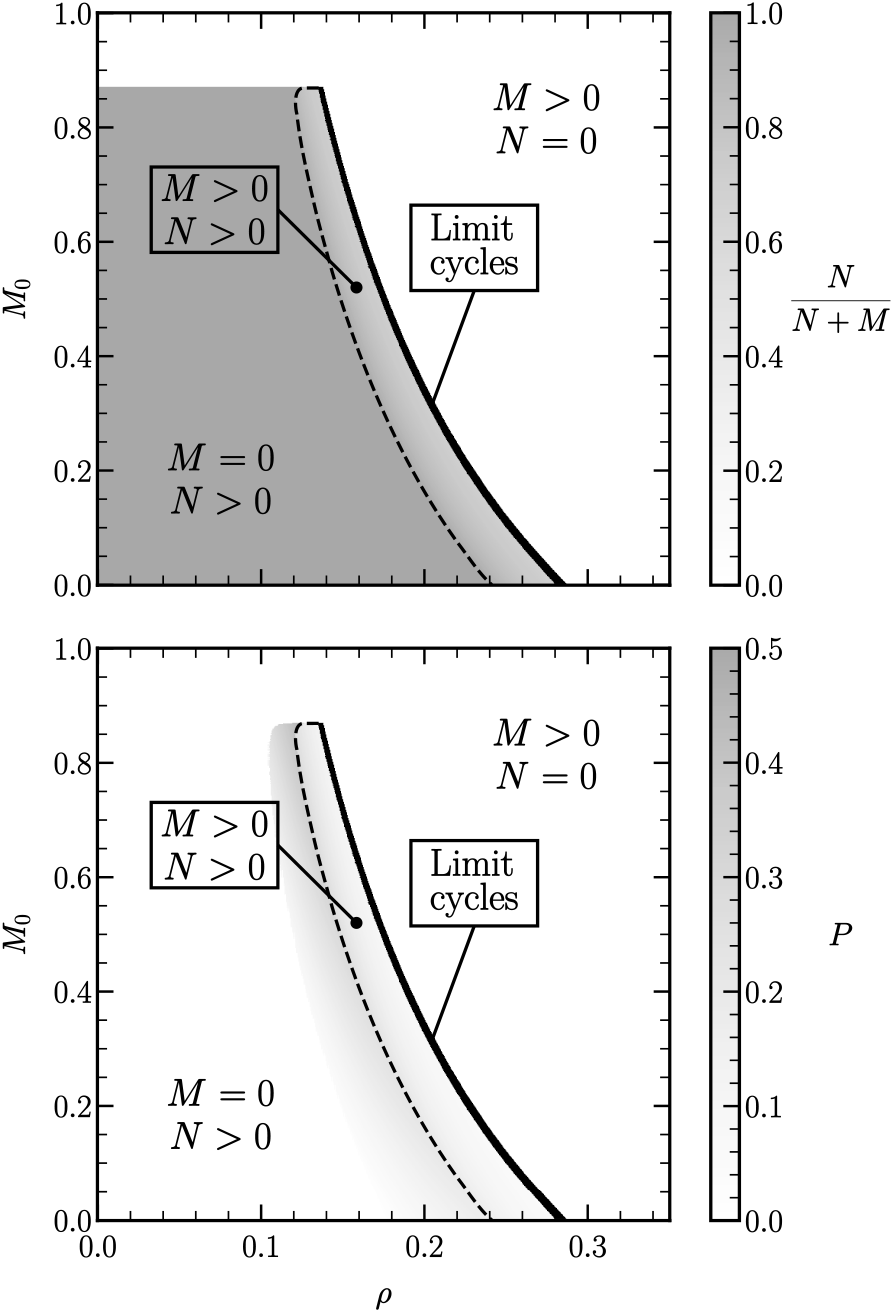
Steady state after invasion by mirror inoculum in a closed system with a predator for the natural counterpart. Increasing predator efficiency (*ρ*) suppresses the natural prey population, opening a niche for mirror-image invasion. The dashed black line outlines the parameter regime where *N, M*, and *P* coexist. The black region marks the transition from this steady-state dominance to a regime where the system no longer settles at a fixed point but instead enters a limit cycle with all three organisms coexisting. Further increasing *ρ* drives a transition to a steady state dominated by the mirror organisms. Parameters: *α* = 1, *β* = 1, *γ* = 0.75, *δ* = *δ*_*p*_ = 0.15, *λ* = 0.1.

**FIG. 5.**
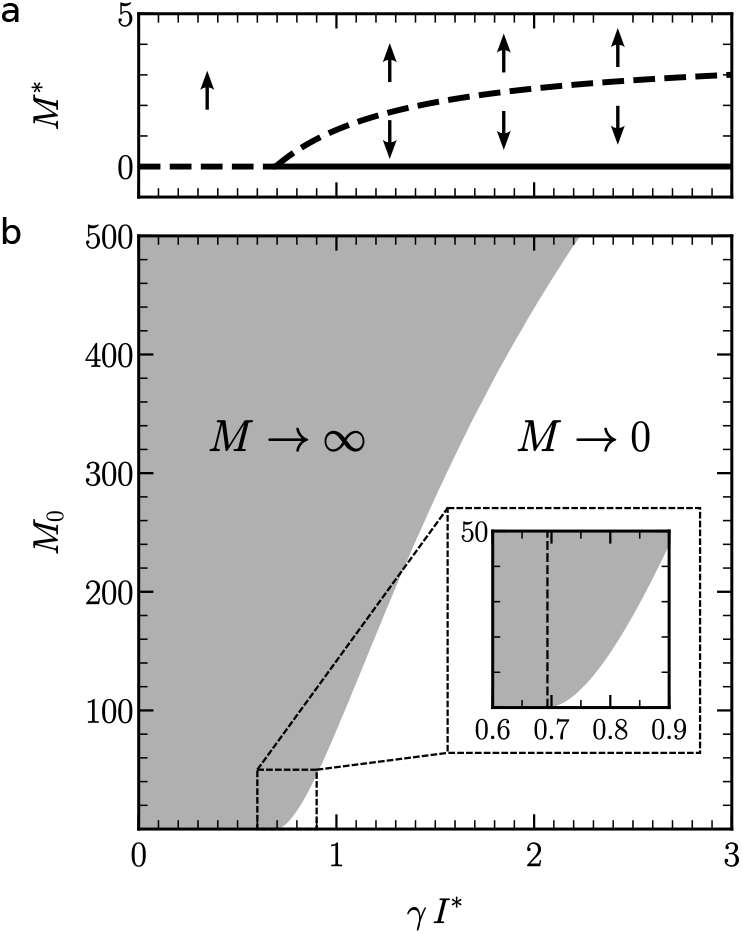
Bifurcation diagram and basins of attraction for the immune response. Panel (a) shows the bifurcation diagram for *M* in system (12) as a function of the clearance rate *γ*. Dashed lines are used to represent unstable fixed points, and solid lines for attractors. At low clearance rates, the immune system cannot effectively recognize or eliminate the mirror organism, and even a small inoculum can escape control. When clearance is sufficiently high, *γ > α δ β*^*−*1^, a saddle–node bifurcation occurs, and the immune system becomes capable of eliminating the threat—provided the initial inoculum is small enough. Panel (b) shows the basin of attraction of the *M* = 0 steady state, computed numerically from initial conditions with *M* = *M*_0_ and macrophages at their steady-state level *I*\* = *β/δ*. Notice that, because the resident macrophage population is initially high, the system proves orders of magnitude more resilient to invasion than the saddle point 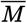 alone would suggest. The inset shows a zoom in the region where the bifurcation takes place (*γ I*\* = *α*). Parameters: *α* = log(2) h^*−*1^, *δ* = log(2)*/*384 h^*−*1^ [46], *β* = 2.7 cells mm^*−*3^ h^*−*1^ [47, 48], *η* = 1 [49]. See the full derivation in SM V. This keeps the units for bacteria interpretable, *M ∝* CFU mm^*−*3^.

In this model, we assume that the immune system recognizes the mirror bacteria, even if some forms of recovery are likely to be impaired. The degree of recognition and elimination is represented by the predation rate (Fig. 5). Generally, the immune system is capable of recognizing foreign or non-self-associated structures that are not strictly dependent on molecular chirality, such as conserved molecular patterns (e.g., glycans, lipopolysaccharides, or physical features of cell surfaces) [50, 51]. This suggests that mirror bacteria would not be completely invisible to innate immune responses, even if specific receptor–ligand interactions are impaired. Note that this dynamical system represents direct entry into susceptible tissue (e.g., via an injury), bypassing physical and chemical barriers. The ingested route poses additional hurdles: it surviving stomach acid and bile acids, then competing with the resident microbiota to establish itself within the gut (which connects us again with the multispecies ecological problem). This scenario could be addressed using the ecosystem dynamics similar to the previous sections.

## DISCUSSION

The biosphere seems locked on one chirality. Could a mirror organism invade it? Invasion dynamics is not reduced to the properties of the invader: its ecological context, from potential competitors to biodiversity, plays an important role in preventing spread [52–54]. An important result from field studies and theoretical models is that biodiversity can act as a firewall to invaders [55– 58]. In this context, our work suggests that the prevailing view that mirror organisms pose an inherent and uncontrollable risk is challenged from an ecological perspective (see Table I). Rather than being a passive environment, the biosphere acts as an active and highly structured system that strongly constrains the establishment and spread of biological novelty, even the extreme case addressed in this work. Our models (a standard approach to invasion dynamics) indicate that the intrinsic nonlinearities associated with resource incompatibility and ecological competition function as an effective form of distributed containment.

These results have direct implications for the framework for biosafety in synthetic biology. Current regulatory approaches emphasize intrinsic molecular or organismal traits [59–65] or rely on institutional protocols to ensure physical isolation [66, 67], while largely treating the environment as passive. This perspective underpins a containment-first paradigm. Our findings challenge this view: viability on the test tube does not necessarily imply establishment. The biosphere acts as an active ecological and biochemical filter that limits the persistence of organisms outside its norms. A more effective framework would therefore shift from organism-centered control to system-level risk assessment, where ecological context is the primary determinant of outcome. In this light, biosafety depends not only on designing constrained organisms but also on maintaining the resilience of ecosystems that naturally restrict their spread. Rather than framing mirror life as a binary choice—prohibit or pursue—the key question is how it would behave within real ecological networks and an evolving biosphere. Addressing this requires moving beyond abstract risk debates toward a grounded system-level understanding of both constraints and possibilities.

Future work should extend this framework to incorporate stochastic effects, population-genetic processes, and heterogeneous communities, as well as explicit evolutionary dynamics. Such extensions are essential to capture rare events, spatial structure, and adaptive responses that could influence invasion outcomes over long timescales. Integrating these dimensions will provide a more complete understanding of how mirror life can behave in realistic biospheric contexts.

## Supporting information

Supplementary Material with extended models.

## ACKNOWLEDGMENTS

We thank the members of the Complex Systems Lab for their valuable discussions and insights. Special thanks to Kevin Esvelt, Richard Lenski, David Relman and Jack Szostak for useful comments and criticisms. R.S. was supported by the AGAUR 2021 SGR 0075 grant, the AEI PID2023-152129NB-I00 grant, and by the Santa Fe Institute. J.P.M. was supported by grant PRE2020-091968, funded by MCIN/AEI (10.13039/501100011033), and co-funded by the ESF through the program “Investing in your future”. VdL is funded by the NYMPHE (HORIZON-CL6-2021-UE 101060625) Contract of the European Union.

## Footnotes

1 By Vieta’s formulas, the product of the roots for a quadratic equation of the generic form *c*_2_*x*^2^ + *c*_1_*x* + *c*_0_ is given by the ratio *c*_0_*/c*_2_. Since *c*_2_ *>* 0 and *c*_0_ *<* 0, the product of the roots is negative, implying that the two values must have opposing signs.

