## Supplementary Material with extended models. for "Ecological constraints to mirror life"

### SUPPLEMENTARY MATERIAL: Ecological constraints to mirror life

Jordi Pla-Mauri,[1,2,*] Victor Maull,[1,2,*] Andrea Tabi,[1,3] Yelyzaveta Shpilkina,[1,2]
Victor de Lorenzo,[4] and Ricard Solé[1,5,2,6,†]
[1]Complex Systems Lab, Universitat Pompeu Fabra, Dr. Aiguader 88, 08003 Barcelona.
[2]Institut de Biologia Evolutiva, CSIC-UPF, Pg. Marítim de la Barceloneta 37, 08003 Barcelona.
[3]Institut de Ciències del Mar, CSIC-ICM, 08003 Barcelona, Spain.
[4]Centro Nacional de Biotecnologia, CSIC, Madrid, Spain.
[5]Institució Catalana de la Recerca i Estudis Avançats (ICREA), Pg. Lluís Companys 23, 08010 Barcelona.
[6]Santa Fe Institute, 1399 Hyde Park Road, Santa Fe, NM 87501, United States.

#### I. AUTOTROPH IN A CLOSED ECOSYSTEM

We consider a well-mixed closed system containing an autotrophic microbial population and a single limiting co-factor (e.g., phosphorous, nitrogen or trace metals, but not a carbon or energy source). The population consists of two strains: a natural resident $N$, which is the established wild type with mortality rate $\delta_n$ and co-factor uptake rate $\alpha_n$; and an invader $M$, a rare mirror-image cell of opposite chirality, differing in both its mortality rate $\delta_m$ and its co-factor uptake rate $\alpha_m$.

Living cells take up the free co-factor $Z$ and incorporate it into catalytic machinery. When cells die, their co-factor content is instantly recycled back into the free pool. The dynamics are therefore

$$\begin{aligned}\frac{dN}{dt} &= N\,(\alpha_n Z - \delta_n),\\ \frac{dM}{dt} &= M\,(\alpha_m Z - \delta_m),\\ \frac{dZ}{dt} &= \delta_n N + \delta_m M - (N\,\alpha_n + M\,\alpha_m)\,Z.\end{aligned} \tag{1}$$

Mass conservation implies that the sum of the derivatives satisfies

$$\frac{dN}{dt} + \frac{dM}{dt} + \frac{dZ}{dt} = 0 \implies Z(t) = Z_0 - N(t) - M(t),$$

so the total co-factor in the system is a conserved quantity, $Z_0$, which can be taken as the initial concentration.

With $M = 0$, a non-trivial equilibrium exists and is globally stable (for $N(0) > 0$) provided $Z_0 > \delta_n/\alpha_n$,

$$N^* = Z_0 - \frac{\delta_n}{\alpha_n}, \qquad Z^* = Z_0 - N^* = \frac{\delta_n}{\alpha_n}.$$

Invasion fitness is the exponential growth rate of the mirror mutant when it is vanishingly rare ($M \to 0$) in the environment set by the resident at equilibrium. Hence

$$r_m^{\text{inv}} = \alpha_m Z^* - \delta_m = \alpha_m \frac{\delta_n}{\alpha_n} - \delta_m = \delta_n \frac{\alpha_m}{\alpha_n} - \delta_m.$$

Mutant invasion succeeds whenever $r_m^{\text{inv}} > 0$, i.e.,

$$\frac{\alpha_m}{\delta_m} > \frac{\alpha_n}{\delta_n}.$$

Thus, in this co-factor-recycling closed ecosystem, selection favors the strain that maximizes the ratio $\alpha/\delta$. A mirror cell with opposite chirality can win even with a lower (or at best equal) growth rate, provided its reduction in mortality from escaping predators and phages outweighs any catalytic handicap.

---

* These two authors contributed equally

This sets up a fundamental trade-off. In the natural ecosystem, cooperation with other organisms can boost the autotroph's replication efficiency, giving the natural strain a higher growth rate ($\alpha_n > \alpha_m$). The mirror cell, by contrast, lacks these cooperative interactions and therefore suffers a catalytic handicap. However, the mirror cell's opposite chirality renders it effectively invisible to phages, while protist predators, and chiral antibiotics will have less effect—all of which might impose strong selective pressure on the natural organism. Consequently, the mirror strain enjoys a substantially lower mortality rate ($\delta_m < \delta_n$). The outcome of invasion thus hinges on whether the mirror cell's survival advantage outweighs its growth disadvantage.

#### II. ANTIBIOTICS AND MICROBIAL WARFARE

Under the assumption of full resource recycling, a closed ecosystem admits the following dynamical description of its biotic steady state:

$$\begin{aligned}
\frac{dN}{dt} &= N\left[\sigma\left(\alpha A + \beta C_n + \gamma C_m\right) - \delta_n\right], \\
\frac{dM}{dt} &= M\left[\varsigma\left(\alpha A + \beta C_m + \gamma C_n\right) - \delta_m\right], \\
\frac{dA}{dt} &= \lambda(\delta_n N + \delta_m M) - \alpha A(\sigma N + \varsigma M), \\
\frac{dC_n}{dt} &= N\left[(1-\lambda)\delta_n - \sigma\beta C_n\right] - \varsigma\gamma M C_n, \\
\frac{dC_m}{dt} &= M\left[(1-\lambda)\delta_m - \varsigma\beta C_m\right] - \sigma\gamma N C_m.
\end{aligned} \tag{2}$$

The multiplicative factors $\sigma, \varsigma \in [0,1]$ modulate the maximum growth rates of the natural and mirror populations, encoding the influence of external growth-limiting agents such as bacteriostatic antibiotics (e.g., chloramphenicol, tetracyclines), which typically target chiral structures including ribosomes and cell wall precursors. The parameters $\delta_n$ and $\delta_m$ represent total per-capita death rates. Bacteriolytic agents ($\beta$-lactams, antimicrobial peptides) increase the death rate through membrane disruption or lysis instead. An implicit form of predation can also be modeled by an increased death rate.

With $M = 0$, the mirror resource $C_m$ decays to zero, and the resident equilibrium gives the steady-state resource concentrations

$$A^* = \frac{\lambda\delta_n}{\alpha\sigma}, \qquad C_n^* = \frac{(1-\lambda)\delta_n}{\sigma\beta}.$$

The resident population $N^*$ is then determined by mass conservation, $\Omega = N^* + A^* + C_n^*$, though its explicit value is not required for the invasion analysis.

The invasion fitness of the mirror variant is its per-capita growth rate when rare, evaluated at this resident state

$$r_m^{\text{inv}} = \varsigma\left(\alpha A^* + \gamma C_n^*\right) - \delta_m = \frac{\varsigma\delta_n}{\sigma}\left[\lambda + \frac{\gamma}{\beta}(1-\lambda)\right] - \delta_m.$$

Setting $r_m^{\text{inv}} > 0$ yields the invasion condition

$$\frac{\varsigma\delta_n}{\sigma\delta_m}\left[\lambda + \frac{\gamma}{\beta}(1-\lambda)\right] > 1.$$

The term in brackets is strictly less than unity whenever $\gamma < \beta$, capturing the mirror organism's inherent handicap: it can only partially utilize the compatible chiral resources the resident ecosystem has accumulated. Successful invasion therefore demands that the mirror compensates with a sufficiently favorable ratio of growth efficiencies to death rates.

Predators, phages, and competing species all accelerate natural population turnover. This raises the steady-state resource concentrations $A^*$ and $C_n^*$ and thereby increases the numerator of the prefactor. Bacteriolytic antibiotics act in the same way. Because these agents evolved against a homochiral world, the mirror's death rate $\delta_m$ may remain largely unaffected, tilting the ratio $\delta_n/\delta_m$ in favor of invasion (see different cases in Figure 1).

Bacteriostatic antibiotics suppress the natural population's ability to convert resources into biomass, lowering $\sigma$ and thus shrinking the denominator of the prefactor—again, often with little impact on the mirror's $\varsigma$. The mirror is constitutively resistant to agents that target L-chiral ribosomes or cell-wall precursors. See Figure 2 for some cases on mirror invasion with different $\sigma$–$\varsigma$ values.

In both scenarios, $r_m^{\text{inv}}$ is driven toward positive values while the chiral penalty stays fixed.

The resident microbiome thus functions as an implicit ecological firewall against invasion, yet external perturbations that selectively weaken the natural side erode the very competitive exclusion that keeps the invasion condition at bay. This yields a counterintuitive outcome: repurposed antibiotics that show some efficacy against mirror organisms in laboratory tests can, when introduced into a microbiome, end up promoting the very invasion they were meant to suppress—by clearing away the natural competitors that would otherwise exclude the mirror from the community.

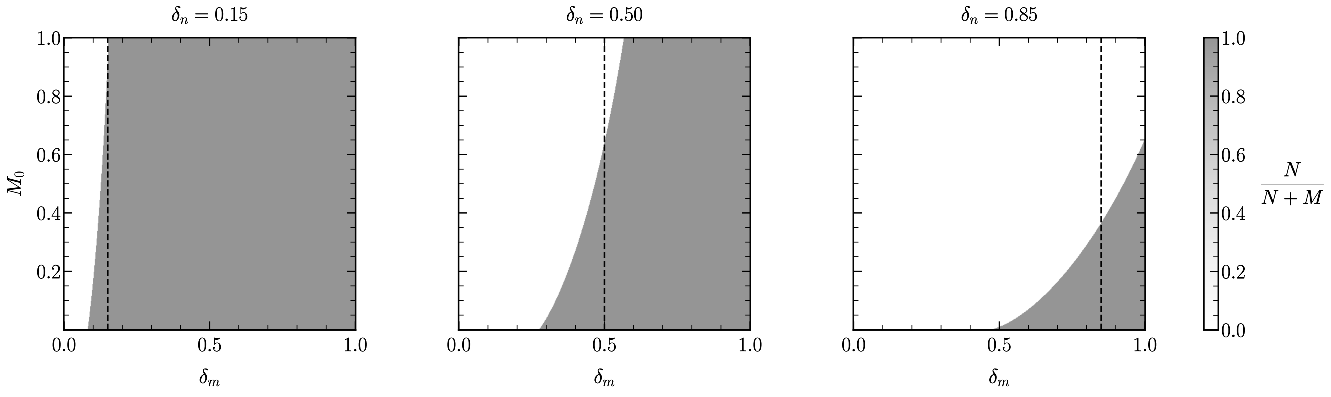


FIG. 1 Heatmaps illustrating mirror invasion dynamics in closed ecosystems under varying death rate asymmetries between natural and mirror populations. Each panel corresponds to a different fixed value of $\delta_n$, varying $\delta_m$ and initial mirror population size $M_0$. The grayscale shading represents the relative abundance at steady state of the natural *versus* mirror population, with dark shades indicating a failed invasion. The vertical dashed line in each panel marks the symmetry point $\delta_m = \delta_n$, where both populations experience identical mortality rates. As shown, when the inoculum size $M_0$ is small, successful invasion by the mirror population occurs only if its death rate is strictly lower than that of the natural population ($\delta_m < \delta_n$). This reflects a competitive advantage conferred by reduced mortality under resource-limited conditions. For larger inocula, however, the mirror population can persist or even dominate even when $\delta_m \geq \delta_n$, due to implicitly adding a large amount of mirror resources when adding mirror biomass. Parameters: $\alpha = \beta = \sigma = \varsigma = 1$, $\gamma = 0.5$, and $\lambda = 0.1$.

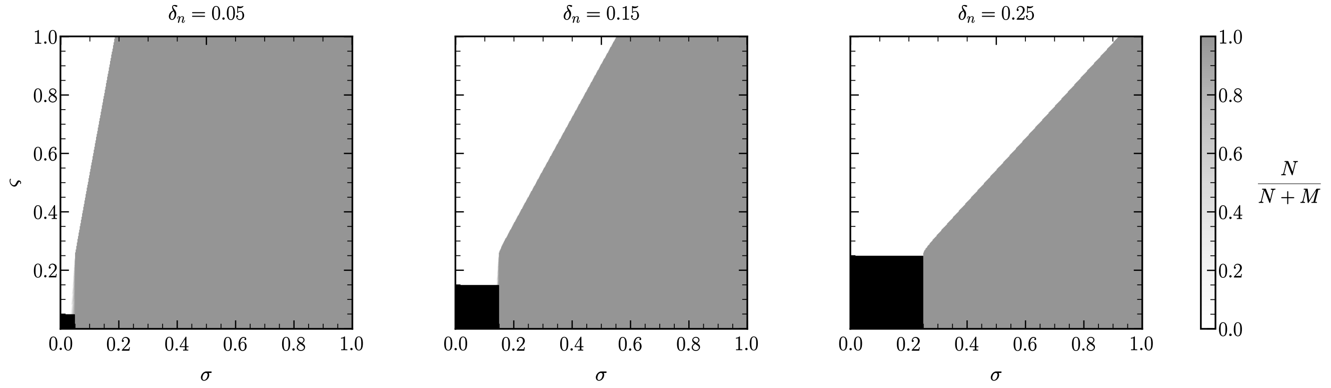


FIG. 2 Heatmaps illustrating mirror invasion dynamics in closed ecosystems under varying growth and death asymmetries between natural and mirror populations after a rare mirror invasor is introduced. Each panel corresponds to a different fixed value of $\delta_n$, with varying $\sigma$ and $\varsigma$ parameters controlling growth suppression of the natural and mirror populations, respectively. The grayscale shading represents the relative abundance at steady state of the natural *versus* mirror population, with dark shades indicating a failed invasion. As $\delta_n$ decreases, successful mirror invasion requires progressively stronger suppression of the natural population, shifting the invasion boundary toward larger $\sigma$ values. Black patches indicate the region where neither population is able to grow due to a higher death rate than its maximum growth rate. Parameters: $\alpha = \beta = 1$, $\gamma = 0.5$, $\delta_m = 0.15$, and $\lambda = 0.1$.

#### III. MIRROR LIFE INVASION IN A MULTISPECIES CHEMOSTAT

We study a microbial ecosystem in which $K$ resident species compete for substitutable resources in a chemostat-like environment. We focus on the invasion potential of a *chiral mirror variant* of a resident organism. We formalize this idea using a dynamical systems approach and show that mirror exclusion persists under coarse-graining.

Here we derive the invasion criterion directly from the *full* consumer–resource model, rather than from the coarse-grained reduction. This provides a more general and formally rigorous foundation for the mirror-exclusion argument. The key idea is standard in invasion analysis: one first identifies the resident equilibrium, then linearizes the complete dynamical system around that equilibrium, and finally examines the eigenvalue associated with perturbations in the mirror-population direction. The sign of that eigenvalue determines whether the mirror lineage can establish when rare.

Let

$$\mathbf{N} = (N_1, \ldots, N_K)$$

be resident populations, and introduce a mirror variant $M$ of a focal resident species. We partition resources into three sectors:

$$\mathbf{R} = (\mathbf{A}, \mathbf{C}_n, \mathbf{C}_m),$$

where: $\mathbf{A}$ is the set of achiral resources, $\mathbf{C}_n$: the natural-handed chiral resources and $\mathbf{C}_m$: mirror-handed chiral resources. Here, the set $\mathbf{A} = (A^{(1)}, A^{(2)}, \ldots)$ contains all resource types that are effectively achiral, or whose uptake and metabolic use do not depend on molecular handedness. These are compounds that can in principle be utilized by both the resident organisms and their hypothetical mirror counterparts without any intrinsic stereochemical mismatch. Examples would include simple inorganic nutrients, achiral carbon sources, or other metabolites for which left–right asymmetry is irrelevant to transport, binding, or catalysis.

The set $\mathbf{C}_n = (C_n^{(1)}, C_n^{(2)}, \ldots)$ contains chiral resource types in the handed form that is compatible with the resident community. These are the molecules naturally produced, transformed, and recycled in an ecosystem built by organisms using the standard biochemical chirality of the residents. Each variable $C_n^{(j)}$ denotes the concentration of the $j$-th natural-handed chiral resource. This class may include amino acids, sugars, metabolites, polymers, breakdown products, or other compounds whose biological utility depends on stereochemical recognition. In real biochemical systems, enzymes and transport machinery are highly stereospecific, so a molecule of the correct handedness can be metabolically valuable whereas its mirror image may be unusable or much less efficiently used.

Finally, the set $\mathbf{C}_m = (C_m^{(1)}, C_m^{(2)}, \ldots)$ contains the mirror-handed analogues of the chiral resources in $\mathbf{C}_n$. These are molecules that would be most compatible with the metabolism of a mirror organism but are typically absent or extremely rare in an ecosystem constructed by naturally handed life. Each variable $C_m^{(\ell)}$ denotes the concentration of the $\ell$-th mirror-handed chiral resource. Biochemically, these are not different in gross chemical composition from the corresponding natural-handed molecules; they differ only in stereochemical configuration. However, that difference is ecologically decisive because molecular recognition, enzyme catalysis, membrane transport, and polymer assembly are all sensitive to chirality.

Using this set of variables, the full multispecies model reads:

$$\frac{dA^{(i)}}{dt} = D(S_A^{(i)} - A^{(i)}) - A^{(i)} \left( \sum_{k=1}^{n} \gamma_{ik} N_k + \gamma_{iM} M \right), \tag{3}$$

$$\frac{dC_n^{(j)}}{dt} = D(S_n^{(j)} - C_n^{(j)}) - C_n^{(j)} \left( \sum_{k=1}^{n} \gamma_{jk} N_k + \gamma_{jM} M \right), \tag{4}$$

$$\frac{dC_m^{(\ell)}}{dt} = D(S_m^{(\ell)} - C_m^{(\ell)}) - C_m^{(\ell)} \left( \sum_{k=1}^{n} \gamma_{\ell k} N_k + \gamma_{\ell M} M \right), \tag{5}$$

together with

$$\frac{dN_k}{dt} = N_k \left( \sum_i \mu_{ki} A^{(i)} + \sum_j \mu_{kj} C_n^{(j)} + \sum_\ell \mu_{k\ell} C_m^{(\ell)} - D \right), \qquad k = 1, \ldots, n, \tag{6}$$

$$\frac{dM}{dt} = M \left( \sum_i \mu_{Mi} A^{(i)} + \sum_j \mu_{Mj} C_n^{(j)} + \sum_\ell \mu_{M\ell} C_m^{(\ell)} - D_M \right). \tag{7}$$

The resident community is composed of the $K$ natural-handed populations $N_1, \ldots, N_K$, while $M$ denotes the mirror variant of one focal resident lineage $N_k$, with $k \in \{1, \ldots, K\}$. To analyze invasion, we first consider the resident ecosystem in the absence of the mirror ($M = 0$). Suppose that the resident system possesses an equilibrium

$$\left(A^{(i)*},\, C_n^{(j)*},\, C_m^{(\ell)*},\, N_k^*,\, 0\right),$$

with at least one surviving focal resident species $N_k^* > 0$.

At this resident equilibrium, the resource balances are

$$0 = D(S_A^{(i)} - A^{(i)*}) - A^{(i)*} \sum_{k=1}^{n} \gamma_{ik} N_k^*, \tag{8}$$

$$0 = D(S_n^{(j)} - C_n^{(j)*}) - C_n^{(j)*} \sum_{k=1}^{n} \gamma_{jk} N_k^*, \tag{9}$$

$$0 = D(S_m^{(\ell)} - C_m^{(\ell)*}) - C_m^{(\ell)*} \sum_{k=1}^{n} \gamma_{\ell k} N_k^*. \tag{10}$$

Similarly, each surviving resident $k$ satisfies

$$\sum_i \mu_{ki} A^{(i)*} + \sum_j \mu_{kj} C_n^{(j)*} + \sum_\ell \mu_{k\ell} C_m^{(\ell)*} = D. \tag{11}$$

These equations define the ecological background generated by the resident community. Biologically, this equilibrium is not a neutral resource state: it is a resource distribution actively maintained by the natural-handed biosystem. In particular, we expect generically that

$$C_n^{(j)*} > 0 \qquad \text{for some } j, \qquad C_m^{(\ell)*} \approx 0 \qquad \text{for most or all } \ell.$$

That is, the resident ecosystem sustains a pool of natural-handed chiral resources, whereas mirror-handed analogues are absent or nearly absent. To linearize the full dynamics, define the full state vector

$$\mathbf{Y} = \left(\{A^{(i)}\}, \{C_n^{(j)}\}, \{C_m^{(\ell)}\}, \{N_k\}, M\right)^\top.$$

The system can be written compactly as

$$\frac{d\mathbf{Y}}{dt} = \mathbf{F}(\mathbf{Y}).$$

The Jacobian matrix at the resident equilibrium is

$$J^* = \left.\frac{\partial \mathbf{F}}{\partial \mathbf{Y}}\right|_{\mathbf{Y}=\mathbf{Y}^*}.$$

Because the state variables include many resources and many resident populations, the full Jacobian is large. However, the invasion problem simplifies because we only need to understand how perturbations in the mirror direction behave when the mirror is initially rare.

Let us write a perturbation about the resident equilibrium as

$$A^{(i)} = A^{(i)*} + \delta A^{(i)}, \qquad C_n^{(j)} = C_n^{(j)*} + \delta C_n^{(j)}, \qquad C_m^{(\ell)} = C_m^{(\ell)*} + \delta C_m^{(\ell)},$$

$$N_k = N_k^* + \delta N_k, \qquad M = 0 + \delta M.$$

The linearized system has the form

$$\frac{d}{dt}\begin{pmatrix} \delta \mathbf{R} \\ \delta \mathbf{N} \\ \delta M \end{pmatrix} = \begin{pmatrix} J_{RR} & J_{RN} & J_{RM} \\ J_{NR} & J_{NN} & J_{NM} \\ J_{MR} & J_{MN} & J_{MM} \end{pmatrix} \begin{pmatrix} \delta \mathbf{R} \\ \delta \mathbf{N} \\ \delta M \end{pmatrix},$$

where $\delta\mathbf{R}$ collects all resource perturbations and $\delta\mathbf{N}$ all resident-population perturbations.

The mirror equation (7) has the form $\frac{dM}{dt} = M\Phi(\mathbf{R})$, where

$$\Phi(\mathbf{R}) = \sum_i \mu_{Mi} A^{(i)} + \sum_j \mu_{Mj} C_n^{(j)} + \sum_\ell \mu_{M\ell} C_m^{(\ell)} - D_M. \tag{12}$$

Since $M$ appears as an overall multiplicative prefactor, when evaluated at the resident equilibrium with $M = 0$, the derivatives of the mirror equation with respect to any resource or resident variable are

$$\frac{\partial}{\partial A^{(i)}} [M\Phi(\mathbf{R})] \bigg|_{M=0} = 0,$$

and similarly

$$\frac{\partial}{\partial C_n^{(j)}} [M\Phi(\mathbf{R})] \bigg|_{M=0} = 0, \qquad \frac{\partial}{\partial C_m^{(\ell)}} [M\Phi(\mathbf{R})] \bigg|_{M=0} = 0, \qquad \frac{\partial}{\partial N_k} [M\Phi(\mathbf{R})] \bigg|_{M=0} = 0.$$

Thus the entire last row of off-diagonal couplings vanishes at invasion:

$$J_{MR} = 0, \qquad J_{MN} = 0.$$

Only the derivative with respect to $M$ itself survives:

$$J_{MM} = \frac{\partial}{\partial M} [M\Phi(\mathbf{R})] \bigg|_{\mathbf{Y}=\mathbf{Y}^*} = \Phi(\mathbf{R}^*). \tag{13}$$

The resulting Jacobian at the resident equilibrium takes the block-upper-triangular form

$$J^* = \begin{pmatrix} J_{\text{res}} & \mathbf{b} \\ \mathbf{0}^\top & r_m^{\text{inv}} \end{pmatrix}, \tag{14}$$

where $J_{\text{res}}$ is the Jacobian of the resident resource–consumer subsystem, $\mathbf{b}$ collects the effect of mirror perturbations on the resident subsystem, and the lower-left block vanishes because a rare mirror does not feed back into its own linearized growth through resident perturbations, giving $r_m^{\text{inv}} = J_{MM}$.

Since a block-upper-triangular matrix has eigenvalues equal to the eigenvalues of its diagonal blocks, the full spectrum of $J^*$ consists of: (a) the eigenvalues of the resident subsystem $J_{\text{res}}$, which determine the internal stability of the resident equilibrium; (b) the single eigenvalue $r_m^{\text{inv}}$, which determines the growth or decay of the mirror invader when rare. This is the formal reason the invasion problem reduces to a single scalar threshold even in the full high-dimensional system. An explicit expression for the invasion eigenvalue can be obtained using (13) and (12), leading to:

$$r_m^{\text{inv}} = \sum_i \mu_{Mi} A^{(i)*} + \sum_j \mu_{Mj} C_n^{(j)*} + \sum_\ell \mu_{M\ell} C_m^{(\ell)*} - D_M. \tag{15}$$

This expression has a direct biological interpretation: it is simply the per-capita growth rate of the mirror lineage evaluated in the resident-generated equilibrium environment. The threshold criterion is immediate:

$$r_m^{\text{inv}} > 0 \implies \text{mirror invasion possible,} \tag{16}$$
$$r_m^{\text{inv}} < 0 \implies \text{mirror excluded,} \tag{17}$$
$$r_m^{\text{inv}} = 0 \implies \text{critical invasion threshold.} \tag{18}$$

Thus invasion is governed by the sign of the mirror's linearized growth rate in the ecological environment produced by the resident community.

In a natural resident ecosystem, the mirror-handed resource pool is expected to be negligible:

$$C_m^{(\ell)*} \approx 0.$$

At the same time, at least some natural-handed resource components are present:

$$C_n^{(j)*} > 0 \qquad \text{for some } j.$$

If the mirror differs from the resident only by chirality and not by demographic parameters, then $D_M = D$. Under these assumptions, (15) reduces approximately to

$$r_m^{\text{inv}} \approx \sum_j (\mu_{Mj} - \mu_{kj}) C_n^{(j)*}. \tag{19}$$

Since each coefficient difference is non-positive and at least one is typically strictly negative, the sum is generically negative:

$$r_m^{\text{inv}} < 0. \tag{20}$$

Thus any sufficiently small introduction of the mirror lineage decays exponentially at first order, and the resident equilibrium is linearly stable against mirror invasion. This result is important because it follows directly from the structure of the complete consumer–resource model and from the chirality asymmetry in uptake coefficients together with the environmentally induced scarcity of mirror-handed resources.

If the mirror has a lower effective loss rate than the resident, so that $D_M < D$, then the last term in (15) becomes positive. The invasion threshold is then

$$\sum_j (\mu_{Mj} - \mu_{kj}) C_n^{(j)*} + \sum_\ell (\mu_{M\ell} - \mu_{k\ell}) C_m^{(\ell)*} + (D - D_M) > 0. \tag{21}$$

Equivalently,

$$D - D_M > \sum_j (\mu_{kj} - \mu_{Mj}) C_n^{(j)*} - \sum_\ell (\mu_{M\ell} - \mu_{k\ell}) C_m^{(\ell)*}. \tag{22}$$

This inequality says that invasion is possible only if the mirror's demographic advantage exceeds its ecological penalty from poor matching to the resident-maintained natural-handed environment. Since the second term on the right-hand side is typically very small, the threshold is usually controlled by the mismatch on the natural-handed resource pool.

The central result of this analysis is that chirality generates an ecological asymmetry that is not merely biochemical, but dynamical and systemic. A mirror organism introduced into an established natural community is not entering a neutral environment. Rather, it enters a resource landscape that has already been constructed, maintained, and stabilized by populations using the opposite molecular handedness. This fact alone strongly constrains the possibility of invasion.

Our derivations show that the appropriate mathematical object for assessing invasion is the mirror growth rate evaluated at the resident equilibrium. In the full consumer–resource system, this quantity appears as the eigenvalue associated with the mirror direction in the Jacobian matrix linearized around the resident-only equilibrium. Because the mirror population enters at vanishing abundance, its initial dynamics are governed by the linear equation

$$\frac{dM}{dt} = r_m^{\text{inv}} M,$$

where $r_m^{\text{inv}}$ is the invasion eigenvalue. The sign of this eigenvalue determines whether the mirror lineage grows or decays when rare. Thus the problem of mirror invasion is reduced to a threshold question in linear stability theory.

A key formal result is that, at the resident equilibrium, the full Jacobian takes a block-triangular form. This happens because the mirror equation contains $M$ as an overall prefactor. As a consequence, when evaluated at $M = 0$, the derivatives of the mirror equation with respect to resident populations and resource variables vanish in the lower-left block of the Jacobian. This means that the eigenvalue governing mirror invasion decouples from the rest of the spectrum and can be identified directly with the mirror's per-capita growth rate in the resident-generated environment. The biological content of this mathematical simplification is important: a rare mirror lineage does not yet have enough abundance to restructure the environment. It must initially survive in whatever ecological conditions the resident community has already created.

This point is crucial, because the resident environment is not chirally symmetric. The natural community consumes, transforms, secretes, and recycles molecules in ways that preferentially maintain the natural-handed resource pool. In our notation, the resource state generated by residents is characterized by the persistence of natural-handed chiral compounds,

$$C_n^{(j)*} > 0 \qquad \text{for some } j,$$

together with the near absence of the mirror-handed analogues,

$$C_m^{(\ell)*} \approx 0.$$

This asymmetry is not imposed externally but arises as an emergent ecological property of the resident biosphere. The natural community does not merely use natural-handed molecules; through its collective metabolism it also maintains an environment in which those molecules remain the dominant biologically relevant chiral forms.

The invasion eigenvalue derived from the full system makes this mechanism explicit. For a mirror analogue of a focal resident species $k$, we obtained

$$r_m^{\text{inv}} = \sum_j (\mu_{Mj} - \mu_{kj}) C_n^{(j)*} + \sum_\ell (\mu_{M\ell} - \mu_{k\ell}) C_m^{(\ell)*} + (D - D_M). \tag{23}$$

This decomposition has a transparent interpretation. The first term measures the mirror's disadvantage on the natural-handed resource pool, the second term measures its potential advantage on the mirror-handed pool, and the third term accounts for any non-chiral demographic difference such as a lower effective dilution or mortality rate.

Under the assumptions appropriate for a genuine mirror analogue, the achiral sector cancels from the comparison because both lineages use achiral resources equally well. Thus achiral resources cannot by themselves rescue the invader. The competition is decided entirely by the chiral sectors of the environment. But here the asymmetry is decisive. The mirror lineage is assumed to be less efficient on the natural-handed pool,

$$\mu_{Mj} \le \mu_{kj},$$

and better matched to the mirror-handed pool,

$$\mu_{M\ell} \ge \mu_{k\ell}.$$

If both resource classes were comparably present, then one could imagine a trade-off between disadvantage in one chiral class and advantage in the other. However, this is not the ecological situation that a rare invader actually encounters. The resident community maintains the first pool but not the second. Therefore the negative contribution to $r_m^{\text{inv}}$ is weighted by appreciable concentrations $C_n^{(j)*}$, whereas the positive contribution is weighted by negligible concentrations $C_m^{(\ell)*}$.

This is the core reason mirror invasion fails. The problem is not simply that mirror organisms are intrinsically inefficient. Rather, they are mismatched to the dominant ecological niche available at the time of introduction. Their preferred resource class is absent because it is not being produced and recycled by the resident biosphere. In this sense, mirror exclusion is a consequence of environmental construction. Established natural life shapes the environment into a state that is selectively favorable to its own chirality and unfavorable to the opposite one.

#### IV. CLOSED ECOSYSTEM WITH MULTIPLE POPULATIONS AND MULTIPLE RESOURCES

We consider a closed ecosystem in which a resident community of $K$ natural-handed microbial species coexists with a finite resource pool of $C$ different resources. All biomass ultimately recycles to the resource pool through death and decomposition. Organisms may also actively convert resources via metabolic pathways including racemases.

We analyze invasion by a chiral mirror variant of a focal resident species $k$.

The resource vector

$$\mathbf{R} = (\mathbf{A}, \mathbf{C}_n, \mathbf{C}_m)^\top \in \mathbb{R}_{\ge 0}^C$$

collects all possible chemical species in the environment that can act as limiting resources. The total resources can be partitioned into three disjoint sets based on chiral properties. Each resource belongs to exactly one of three distinct categories: achiral resources

$$\mathbf{A} = (A^{(1)}, A^{(2)}, \dots)^\top,$$

which are symmetric molecules, inorganic nutrients, or simple organic acids that possess no handedness or exhibit biological activity independent of chirality; natural-handed chiral resources

$$\mathbf{C}_n = (C_n^{(1)}, C_n^{(2)}, \dots)^\top,$$

which consist of chiral molecules in the stereochemical configuration compatible with natural-handed life; and mirror-handed chiral resources

$$\mathbf{C}_m = (C_m^{(1)}, C_m^{(2)}, \dots)^\top,$$

which comprise the corresponding chiral molecules in the opposite stereochemical configuration.

The resident biomass vector,

$$\mathbf{N} = (N_1, \dots, N_K)^\top \in \mathbb{R}^K_{\geq 0},$$

collects the population sizes of all $K$ resident species. An additional mirror population $M \in \mathbb{R}_{\geq 0}$ denotes the invader population. Together, the full system state is $(\mathbf{R}, \mathbf{N}, M)$.

Each consumer $X$ possesses a growth function $F_X(\mathbf{R})$. The function is assumed to be smooth, non-decreasing in each argument, and satisfies $F_X(\mathbf{0}) = 0$. Moreover, to cleanly partition growth contributions by chirality, we assume the growth function is additive across different resource categories

$$F_X(\mathbf{R}) = F_X\left((\mathbf{A}, \mathbf{0}, \mathbf{0})^\top\right) + F_X\left((\mathbf{0}, \mathbf{C}_n, \mathbf{0})^\top\right) + F_X\left((\mathbf{0}, \mathbf{0}, \mathbf{C}_m)^\top\right),$$

which assumes that growth contributions from achiral, natural-handed, and mirror-handed resources are metabolically independent (i.e., no cross-chiral synergy or inhibition). The monotonicity condition

$$\frac{\partial F_X}{\partial \mathbf{R}} \geq \mathbf{0}$$

reflects that adding more of any resource never reduces growth; resources are substitutable but never inhibitory at the concentrations considered. However, because growth may be constrained by a different limiting resource, adding more of a non-limiting resource can result in zero net increase in growth, satisfying the inequality at zero.

The growth function $F_X(\mathbf{R})$ can be supported only on a specific subset of resources. For any resource that does not contribute to growth, the corresponding partial derivative is identically zero regardless of concentration; where growth is supported, the partial derivative is strictly positive in some region of resource space.

Resident species naturally thrive on achiral and natural-handed resources, which constitute their primary metabolic base, but can also exploit a limited subset of mirror-handed resources with low efficiency via native racemases or promiscuous enzymes. For a given resident species $k$, growth exhibits non-zero partial derivatives with respect to some achiral resources and natural-handed resources, possibly alongside a weak sensitivity to a small subset of mirror-handed resources. This metabolic asymmetry is encoded by the componentwise inequality

$$\frac{\partial F_k}{\partial \mathbf{C}_m} \leq \frac{\partial F_k}{\partial \mathbf{C}_n},$$

where equality holds only in the absence of any cross-chiral conversion pathway.

The mirror invader $M$ is a synthetic variant of the focal resident species $k$ with completely inverted macromolecular chirality. In extreme cases, it may include engineered sensitivity to additional natural-handed resources which the natural counterpart cannot access in mirror form.

Because inverting stereochemistry does not alter interactions with achiral molecules, the focal resident and its mirror share identical growth responses to achiral resources,

$$\frac{\partial F_k}{\partial \mathbf{A}} = \frac{\partial F_M}{\partial \mathbf{A}}.$$

The resident biomass vector and the mirror population follow

$$\frac{d\mathbf{N}}{dt} = \mathbf{N} \odot (F(\mathbf{R}) - \boldsymbol{\delta}), \qquad \frac{dM}{dt} = M\left(F_M(\mathbf{R}) - \delta_M\right),$$

where the vector of growth functions

$$F(\mathbf{R}) = (F_1(\mathbf{R}), \dots, F_K(\mathbf{R}))^\top,$$

the vector of resident death rates

$$\boldsymbol{\delta} = (\delta_1, \dots, \delta_K)^\top,$$

and $\odot$ denotes the elementwise product. Loss rates $\boldsymbol{\delta}$ and $\delta_M$ are assumed constant given the subsequent analysis is focused at steady state. Any reduction $\delta_M < \delta_k$ reflects the mirror's escape from chirally specific antagonistic interactions such as antibiotics, bacteriocins, or phages.

Let the consumption matrix $\mathbf{Q}(\mathbf{R}) \in \mathbb{R}^{C\times(K+1)}$ collect the per-capita consumption vectors of all consumers as columns, where column $X$ corresponds to the per-capita consumption vector of consumer $X$ and is denoted $\mathbf{Q}_X(\mathbf{R})$. We assume a unitary biomass yield to enforce mass conservation,

$$\mathbf{1}^\top \mathbf{Q}_X(\mathbf{R}) = F_X(\mathbf{R}),$$

which states that total per-capita resource consumption equals the growth rate. The allocation of this total consumption across individual resources is determined by the organism's metabolic strategy.

Decomposition is strictly chirality-conserving. The recycling matrix $\mathbf{D} \in \mathbb{R}^{C\times(K+1)}$ collects the recycling vectors of all consumers as columns. Columns $\mathbf{D}_X$, corresponding to residents $X \in \mathbf{N}$, have support restricted to $(\mathbf{A}, \mathbf{C}_n, \mathbf{0})^\top$. Column $\mathbf{D}_M$, corresponding to the mirror invader $M$, has support restricted to $(\mathbf{A}, \mathbf{0}, \mathbf{C}_m)^\top$. The whole matrix satisfies

$$\mathbf{1}^\top \mathbf{D}_X = 1, \qquad X \in \mathbf{N} \cup \{M\}.$$

Active metabolic conversion is encoded in the per-capita rate matrix $\mathbf{K}_X \in \mathbb{R}^{C\times C}$ for each consumer population $X$, with entries $(\mathbf{K}_X)_{R,R'} = \kappa(R, R', X)$. The matrix satisfies

$$\mathbf{1}^\top \mathbf{K}_X = \mathbf{0}^\top, \qquad X \in \mathbf{N} \cup \{M\},$$

to ensure that metabolic conversion merely redistributes mass among resource categories without creating or destroying it; off-diagonal blocks coupling $\mathbf{C}_n$ and $\mathbf{C}_m$ are permitted and encode racemase activity.

The resource dynamics are

$$\frac{d\mathbf{R}}{dt} = \sum_X X\left[\delta_X \mathbf{D}_X - \mathbf{Q}_X(\mathbf{R}) + \mathbf{K}_X \mathbf{R}\right].$$

Total mass conservation follows as

$$\frac{d}{dt}\left(\mathbf{1}^\top \mathbf{R} + \mathbf{1}^\top \mathbf{N} + M\right) = 0,$$

which holds since $\mathbf{1}^\top \mathbf{D}_X = 1$ ensures each unit of biomass lost to death is fully returned to the resource pool, and $\mathbf{1}^\top \mathbf{K}_X = \mathbf{0}^\top$ ensures metabolic conversion merely redistributes mass among resource categories without creating or destroying it.

Without the mirror organism present ($M = 0$), the resident community is assumed to reach a stable equilibrium $(\mathbf{R}^*, \mathbf{N}^*, 0)$ with at least resident $N_k^* > 0$. The equilibrium conditions are

$$F(\mathbf{R}^*) = \boldsymbol{\delta}, \qquad \sum_{k=1}^{K} N_k^* \left[\delta_k \mathbf{D}_k - \mathbf{Q}_k(\mathbf{R}^*) + \mathbf{K}_k \mathbf{R}^*\right] = \mathbf{0}.$$

For mirror-handed resources $\mathbf{C}_m$, the columns $\mathbf{D}_k$ of all natural residents have zero support on $\mathbf{C}_m$, so mirror-handed resources are produced at steady state only through weak racemase-mediated conversion from natural-handed precursors. The equilibrium concentrations $\mathbf{C}_m^*$ are therefore assumed to be small. They may not completely vanish, but are maintained at low levels by the balance of weak production and weak consumption by resident organisms.

The invasion fitness is

$$r_m^{\text{inv}} = F_M(\mathbf{R}^*) - \delta_M. \tag{24}$$

To relate this to the focal resident, we use its equilibrium condition $F_k(\mathbf{R}^*) - \delta_k = 0$ to evaluate the conditions for neutral invasion

$$r_m^{\text{inv}} = (\delta_k - \delta_M) - (F_k(\mathbf{R}^*) - F_M(\mathbf{R}^*)). \tag{25}$$

Partitioning growth by chirality, and using the assumption of achiral symmetry, we obtain

$$r_m^{\text{inv}} = (\delta_k - \delta_M) - \Delta_{\mathbf{C}_n} + \Delta_{\mathbf{C}_m}, \tag{26}$$

where $\Delta_{\mathbf{C}_n}$ is the mirror's growth deficit on natural-handed resources and $\Delta_{\mathbf{C}_m}$ is its advantage on mirror-handed resources, defined as

$$\Delta_{\mathbf{C}_n} = F_k(\mathbf{R}_n^*) - F_M(\mathbf{R}_n^*) \geq 0, \qquad \mathbf{R}_n^* = (\mathbf{0}, \mathbf{C}_n^*, \mathbf{0})^\top,$$
$$\Delta_{\mathbf{C}_m} = F_M(\mathbf{R}_m^*) - F_k(\mathbf{R}_m^*) \geq 0, \qquad \mathbf{R}_m^* = (\mathbf{0}, \mathbf{0}, \mathbf{C}_m^*)^\top.$$

Both organisms share identical responses to achiral resources, so the achiral growth contributions cancel exactly.

Successful invasion requires

$$r_m^{\text{inv}} > 0 \implies \delta_k - \delta_M > \Delta_{\mathbf{C}_n} - \Delta_{\mathbf{C}_m}. \tag{27}$$

In natural ecosystems, mirror-handed resources are rare. Abiotic sources are negligible in most settings, and resident organisms produce them only as sparse metabolic byproducts—such as D-amino acids in peptidoglycan or secreted signals—keeping environmental concentrations at trace levels. Consequently, assuming $C_m^{(\ell)*} \approx 0$ and thus $\Delta_{\mathbf{C}_m} \approx 0$, the invasion criterion simplifies to

$$\delta_k - \delta_M > \Delta_{\mathbf{C}_n}. \tag{28}$$

Thus, in order to succeed, a mirror organism must gain more from evading chiral-specific mortality factors than it loses in its ability to exploit the natural-handed resources that dominate the ecosystem. In the absence of an abundant mirror-handed resource base, the mortality advantage represents the sole invasion driver, and its magnitude must be substantial given that $\Delta_{\mathbf{C}_n}$ encompasses the full growth penalty on the resources that sustain the resident community.

#### V. UNIT DERIVATION FOR IMMUNE RESPOSE

##### A. Bacterial Growth Rate

The bacterial growth rate is approximated to

$$\alpha \approx \frac{\log(2)}{1\,\text{h}} \approx 0.693\,\text{h}^{-1},$$

which directly reflects a fast standard doubling time of 1 h during exponential growth under suitable but not-perfect conditions.

##### B. Macrophage Natural Clearance rate

For the macrophage population, the natural clearance rate is approximated as

$$\delta \approx \frac{\log(2)}{384\,\text{h}} \approx 1.8 \times 10^{-3}\,\text{h}^{-1}.$$

This value is derived from an estimated macrophage tissue half-life of 16 days (384 h) reported by Parwaresch *et al.* [1], assuming a first-order decay process

$$t_{1/2} = \frac{\log(2)}{\delta}.$$

##### C. Macrophage Recruitment Rate

The macrophage recruitment rate is approximated to be

$$\beta \approx 2.7\,\text{macrophages}\,\text{mm}^{-3}\,\text{h}^{-1}.$$

This is derived by taking the baseline resident population of $3 \times 10^6$ macrophages in the mouse peritoneal cavity reported by Iyengar *et al.* [2] and distributing it across a physiological fluid volume of 2 ml (2000 $\text{mm}^3$) as established by Zhu *et al.* [3]. Multiplying this baseline density (1500 macrophages $\text{mm}^{-3}$) by the natural clearance rate $\delta$ yields the steady-state recruitment flux required to maintain the resident population in the absence of infection.

##### D. Macrophage Effective Capacity

The reciprocal of the maximum carrying capacity per phagocyte,

$$\eta \approx 1\,\text{macrophages}\,\text{bacteria}^{-1}.$$

This corresponds to an effective saturation threshold of approximately 1 bacteria per macrophage, a reduced version of the intracellular capacity bounds in a lifetime observed by Lesbats *et al.* [4]. This lower value is assumed due to the fact of possible toxicity of the ingested mirrored components for the macrophage.

##### E. Phagocytic Clearance Rate

Pesanti [5] measured phagocytosis of *S. aureus* by mouse peritoneal macrophages *in vitro* using Lineweaver–Burk analysis, reporting uptake rate $V$ in $\text{CFU}\,\text{mg}^{-1}\,\text{min}^{-1}$ of monolayer protein and bacterial concentration $S$ in $\text{CFU}\,\text{ml}^{-1}$.

Fitting the control data from Figure 1 yields $V_{\text{max}} \approx 1.7 \times 10^8\,\text{CFU}\,\text{mg}^{-1}\,\text{min}^{-1}$ and $K_m \approx 7.8 \times 10^7\,\text{CFU}\,\text{ml}^{-1}$.

Because bacterial concentrations in our simulations are assumed to be well below $K_m$, uptake is approximately linear in $S$, and the Michaelis–Menten expression reduces to a first-order uptake rate proportional to $V_{\text{max}}/K_m$.

To convert this quantity into model units ($\text{mm}^3\,\text{macrophage}^{-1}\,\text{h}^{-1}$), we convert $V_{\text{max}}$ from a per-mg-protein basis to a per-cell basis using an estimated macrophage protein content of approximately 300 pg per cell [6]. Given the approximately $3 \times 10^5$ macrophages per monolayer reported in the paper, each monolayer contains roughly 0.09 mg protein, corresponding to approximately $3.3 \times 10^6\,\text{macrophages}\,\text{mg}^{-1}$ protein.

This yields a per-cell $V_{\text{max}}$ of approximately $3.1 \times 10^3\,\text{CFU}\,\text{macrophage}^{-1}\,\text{h}^{-1}$ and a low-density predation coefficient of

$$\gamma \approx \frac{V_{\text{max}}}{K_m} \approx 0.04\,\text{mm}^3\,\text{macrophages}^{-1}\,\text{h}^{-1},$$

under bacteria-limited conditions.

Using the previously established parameter values, the threshold for the critical predation coefficient is given by

$$\gamma_c = \frac{\alpha}{\beta}\delta \approx \frac{\log(2)}{2.7}\frac{\log(2)}{384} \approx 4.6 \times 10^{-4}.$$

The ratio with respect the approximated value for natural organisms yields

$$\frac{\gamma}{\gamma_c} \approx \frac{0.04}{4.6 \times 10^{-4}} \approx 87.$$

So the projected clearance rate of the macrophages exceeds the saddle–node bifurcation threshold by nearly two orders of magnitude. Within this parameter regime, tissue-resident macrophages are expected to robustly stabilize the bacteria-free equilibrium state, provided the initial inocula remain sufficiently small. Thus, for a mirror invasion to initially win, the macrophage clearance efficiency would need to be reduced by almost two orders of magnitude to fall below this critical threshold.
